# BACE1 – but not BACE2 – function is critical for metabolic disorders induced by high-fat diets in C57BL/6N mice

**DOI:** 10.1101/2021.12.20.473491

**Authors:** Thomas W. Rosahl, Lynn A. Hyde, Patrick T. Reilly, Marie-France Champy, Kirstine J. Belongie, Benoit Petit-Demouliere, Tania Sorg, Hugues Jacobs, Robert Terry, Jack D. Scott, Jared N. Cumming, Eric M. Parker, Matthew E. Kennedy

**Affiliations:** Merck & Co., Inc., Kenilworth, NJ, USA; CELPHEDIA, PHENOMIN, Institut Clinique de la Souris (ICS), Illkirch, France; Centre National de la Recherche Scientifique, UMR7104, Illkirch, France; Institut National de la Santé et de la Recherche Médicale, U964, Illkirch, France; Université de Strasbourg, 1 rue Laurent Fries, 67404 Illkirch, France; Merck & Co., Inc., Boston, MA, USA

**Author notes:** Corresponding author: Matthew E. Kennedy, Merck & Co., Inc., Boston, MA 02115, USA.

**Keywords:** Alzheimer’s disease, BACE (β-secretase), diabetes, metabolism, preclinical, TMEM27

## Abstract

**Aims/hypothesis:** Beta-site amyloid precursor protein-cleaving enzyme 1 (BACE1) is required for the production of toxic amyloid peptides and is highly expressed in the brain, but also to a lesser extent in major peripheral organs such as muscle and liver. In contrast, BACE2 is mainly expressed in peripheral tissues and is enriched in pancreatic beta cells, where it regulates beta-cell function and mass. Previous reports demonstrated that loss of BACE1 function decreases body weight, protects against diet-induced obesity and enhances insulin sensitivity in mice, whereas mice lacking *Bace2* exhibit reduced blood glucose levels, improved intraperitoneal glucose tolerance and increased beta-cell mass. Impaired glucose homeostasis and insulin resistance are hallmarks of type 2 diabetes and have been implicated in Alzheimer’s disease. Therefore, we tested the contribution of the individual BACE isoforms to those metabolic phenotypes by placing *Bace1* knockout (KO), *Bace2* KO, *Bace1/2* double knockout (dKO) and wild-type (WT) mice on a high-fat high-cholesterol diet (HFD) for 16 weeks.

**Methods:** *Bace1* KO (*n* = 18), *Bace2* KO (*n* = 18), *Bace1/2* dKO (*n* = 18) and WT C57BL/6N mice (*n* = 54) were fed a HFD for 16 weeks (age 9–25 weeks). Body composition was measured before initiation of the HFD and after 11 weeks of HFD. Oral glucose tolerance and insulin sensitivity tests were performed after 12 and 13 weeks of HFD, respectively, and full blood chemistry was analyzed after 16 weeks of HFD. The effects of subchronic BACE1/2 inhibition were assessed by administration of 10 mg/kg/day of the dual BACE1/2 inhibitor MBi-3 in a HFD fed to C57BL/6N mice for 3 weeks.

**Results:** *Bace1* KO and *Bace1/2* dKO mice showed decreased body weight and improved glucose tolerance and insulin resistance vs. WT mice. Conversely, *Bace2* KO mice did not show any significant differences in body weight, glucose tolerance or insulin resistance under our experimental conditions. Finally, subchronic MBi-3–mediated BACE1/2 inhibition in mice in conjunction with a HFD resulted in a modest improvement of glucose tolerance.

**Conclusions/interpretation:** Our data indicate that lack of BACE1 – but not BACE2 – function contributes mainly to the metabolic phenotypic changes observed in *Bace1/2* dKO mice, suggesting that inhibition of BACE1 has the greater role (vs. BACE2) in any potential improvements in metabolic homeostasis.

**HIGHLIGHTS:**

- Insulin resistance may develop in the brains of patients with Alzheimer’s disease (83/85 characters)
- BACE1 and BACE2 may play a role in glucose homeostasis and insulin sensitivity (80/85 characters)
- Body weight in mice decreased with *Bace1* KO and *Bace1/2* KO but not *Bace2* KO alone (83/85 characters)
- *Bace1* and *Bace1/2,* but not *Bace2,* KO improved glucose tolerance/insulin resistance (84/85 characters)
- Improved metabolic homeostasis may follow loss of BACE1 rather than BACE 2 activity (85/85 characters)

## 1. Introduction

Alzheimer’s disease (AD) is triggered by the abnormal aggregation and accumulation of amyloid beta (Aβ) peptides into toxic oligomers and insoluble extracellular amyloid plaques that, in turn, enhance the formation and spread of neurofibrillary tangles of hyperphosphorylated tau and mobilization of disease-associated microglia [1]. Over time, this leads to synaptic loss and, ultimately, neuronal death in brain regions essential for learning and memory [2–4].

The Aβ peptide is derived from the amyloid precursor protein (APP) via successive cleavage by the beta-site APP-cleaving enzyme 1 (BACE1; beta-secretase 1)—which is predominantly expressed in the brain and pancreas, although beta-secretase enzymatic activity is primarily detected in brain [5, 6]—and gamma-secretases [7]. Consequently, there has been significant investigation into BACE1 inhibitors, with the aim of lowering cerebrospinal fluid (CSF) Aβ and reducing brain amyloid plaque pathology. Several non-selective inhibitors of both BACE1 and BACE2, a close homologue of BACE1, have been shown to lower CSF Aβ levels in preclinical models [8–13], healthy volunteers [9, 13] and patients with AD [8, 9, 14–16]. BACE2 is expressed mainly in the periphery, particularly in pancreatic beta-cells, and is not required for Aβ production [17–19].

BACE1 and BACE2 may play a role in glucose homeostasis and insulin sensitivity, which are hallmarks of type 2 diabetes [19–23] alongside insulin resistance in peripheral tissues [24].

There is also accumulating evidence of insulin resistance developing in the brains of patients with AD [25, 26]. Thus, it is of interest to understand the impact that non-selective BACE inhibitors may have beyond lowering Aβ levels. Germline loss of BACE1 in mice is associated with increased uncoupled respiration and metabolic inefficiency, decreased body weight, protection against diet-induced obesity and enhanced insulin sensitivity [21], while overexpression of BACE1 in mouse neurons induced severe impairment of systemic glucose homeostasis and insulin sensitivity, brain inflammation, and cognitive impairment [22]. Meanwhile, BACE2 is the major sheddase of the plasma membrane protein, TMEM27 [20], which has growth-promoting and insulin-stimulating activity in pancreatic beta-cells [27] and regulates amino acid uptake in the kidney [28]. Inhibition or loss of BACE2 function has been shown to improve control of glucose homeostasis via increased insulin levels and increased beta-cell number without a change in body weight [20].

In this report, we investigated the individual contributions of BACE1 and BACE2 on metabolic phenotypes in control (wild-type [WT]) mice and in mice lacking *Bace1*, *Bace2* or both *Bace1/2* after consumption of a high-fat high-cholesterol diet (HFD) for 16 weeks. In addition, we compared the effects of life-long loss of *Bace1/2* vs. pharmacological inhibition of BACE1/2 in adulthood on similar metabolic endpoints.

## 2. Methods

### 2.1. Ethics statement

All experiments were carried out in accordance with the European Communities Council Directive of November 24, 1986. Dr Yann Herault, as the principal investigator in this study, was granted the approval to perform the reported experiments under protocol numbers 2012-034 and 2014-002. The experimental procedures were authorized by the French Ministry of Research Committee C2EA-17.

The care of animals and experimental protocols applied in this manuscript were reviewed and approved by the MSD Experimental Animal Care and Use Committee.

### 2.2. Animals

All mice used in these studies were on a C57BL/6N background. *Bace1* knockout (KO) mice were generated by Xenogen Biosciences (now Taconic Biosciences Inc., Albany, NY, USA) and *Bace2* KO mice by Lexicon Genetics Inc. (The Woodlands, TX, USA), on behalf of Merck & Co., Inc., Kenilworth, NJ, USA. Further details of the generation of *Bace1* KO, *Bace2* KO and *Bace1/2* double KO (dKO) mice is given in the electronic supplementary material (ESM; Supplementary Methods and Fig. S1).

### 2.3. *BACE1*, *BACE2* and *Tmem27* expression plasmids

The mouse *Tmem27* gene (NM_020626) was expressed from the expression vector pCMV6- Entry, with a myc-FLAG epitope tag on the C-terminus (MR223526, Origene Technologies, Rockville, MD, USA). The human full-length *BACE1* or *BACE2* cDNAs were expressed from the pcDNA3.1/myc-His expression vector (Invitrogen, Waltham, MA, USA).

### 2.4. Primary cultures and cell lines

Medium, serum, and supplements for maintenance of cells were obtained from Invitrogen. HEK293 cells were maintained in Dulbecco’s modified Eagle’s medium (DMEM) supplemented with 10% fetal bovine serum (FBS). HEK293-APP clone 46 cells were derived from HEK293 cells through stable transfection of the human *APPswe* gene and maintained as described previously [29]. Beta-TC-6 cells (CRL-11506, ATCC, Manassas, VA, USA) were maintained in DMEM supplemented with 15% FBS and 1 mM sodium pyruvate. Primary pancreatic islets were isolated as described previously [20] and maintained in Roswell Park Memorial Institute medium supplemented with 10% FBS.

### 2.5. Expression of BACE1 and BACE2 in *Bace2* KO mouse tissue and determination of **A**β **levels**

The preparation of membrane protein samples is described in the ESM. Samples (10 μg) were resolved on 4–12% NuPage Bis-Tris gels (Invitrogen), transferred to nitrocellulose membranes and probed with antibodies against the BACE2 C-terminus (sc-271212, Santa Cruz Biotechnology, Dallas, TX, USA), BACE1 N-terminus (PA5-14879, Invitrogen), or Na+/K+- ATPase (Thermo Fisher Scientific, Waltham, MA, USA). Proteins were detected using electrochemiluminescence and scanned on an Odyssey Fc imager (Li-Cor, Lincoln, NE, USA). Dosing with BACE inhibitors MBi-1, MBi-2 and MBi-7 (ESM Fig. S2), and collection of mouse cortical tissue and plasma for Aβ 40 evaluations were performed as described previously [9].

### 2.6. TMEM27 cleavage in HEK293 cells and primary islets and inhibition by MBi-2

HEK293-APP-Swedish/London cells (obtained from ATCC and stably transfected with human APP cDNA containing the Swedish and London familial AD mutations) were plated in plates 1 day before transfection with a total of 2 μg of DNA (1 μg mouse pCMV6-Tmem27-Flag and 1 μg empty vector or pCDNA3.1-BACE2-myc) and 5 μL Lipofectamine 2000 (Invitrogen).

One day post-transfection, the medium was replaced with 1 mL of medium containing 0.2% dimethyl sulfoxide (DMSO; vehicle) or various concentrations of MBi-2 in 0.2% DMSO. The following day, the conditioned medium was collected, the cells washed with phosphate-buffered saline, and lysed with ice-cold radioimmunoprecipitation assay buffer containing a cocktail of protease and phosphatase inhibitors (HALT, Thermo Fisher Scientific). The lysate was sonicated (2 min), then insoluble material pelleted (20,000 × *g*, 15 min, 4°C). Primary mouse islet lysates were prepared from approximately 100 islets as described above. Protein concentrations of cell lysates were determined by Pierce BCA protein assay (Thermo Fisher Scientific). Equal amounts of cell-lysate protein and equal volumes of supernatant were resolved on 12% NuPage Bis-Tris gels and transferred to nitrocellulose membranes. To assess TMEM27 cleavage, blots were probed with antibodies against the extracellular domain of TMEM27 (LS-C125384, LifeSpan BioSciences, Inc., Seattle, WA, USA) or anti-M2 (FLAG) antibody for full-length TMEM27. APP processing was monitored in the supernatant fraction using an in-house rabbit polyclonal antibody (Research Genetics, Inc, Huntsville, AL, USA) against the C-terminal Swedish mutation neoepitope on soluble APPβ that is formed by BACE1 cleavage. Proteins were detected using IRDye secondary antibodies (Li-Cor) and scanned on an Odyssey Fc imaging system (Li-Cor).

### 2.7. Histological evaluation of islets in *Bace2* KO mice

Pancreata were fixed in 10% formalin, embedded in paraffin and sectioned (5 µm). Deparaffinized sections were blocked in 5% normal goat serum followed by overnight incubation with primary antibodies at 4°C: anti-BACE2 antibody (Santa Cruz; 1:1000), anti-insulin (Invitrogen; 1:100), anti-glucagon (Invitrogen, 1:100) and anti-Ki-67 (BD Biosciences, San Jose, CA, USA; 1:50). Anti-Ki-67 detection required antigen retrieval by boiling in 0.01 M Na-citrate pH 6.0. Slides were scanned on an Aperio ScanScope FL (Leica Microsystems GmbH, Wetzlar, Germany) and images were captured and analyzed using ImageScope v11 and Spectrum v11 (Leica Microsystems) to count islet numbers, islet area (area occupied by insulin- and glucagon-positive cells), nuclei and Ki-67-positive cells.

### 2.8. Metabolic assessment of the *Bace* mutant mice

Metabolic studies were carried out on three separate cohorts of KO mice (*Bace1* KO, *Bace2* KO and *Bace1/2* dKO). Each cohort consisted of 18 mutant mice (nine males, nine females) and 18 WT littermate controls (nine males, nine females). Due to breeding constraints, these cohorts were assessed separately under the same regimen of challenge and tests. Upon arrival, mice were housed three per cage and fed with a standard control chow diet (CD) (D04, Safe, Augy, France). At the age of 9 weeks, the diet was switched to a HFD (RD 12492, 60% kcal fat; Research Diets, New Brunswick, NJ, USA) for 16 weeks.

#### 2.8.1. Body weight and fat deposition

Body weight was recorded once weekly from 5 to 20 weeks of age. To determine fat deposition before and after HFD challenge, body composition (fat tissue, lean tissue and free body fluid) was evaluated by quantitative nuclear magnetic resonance (qNMR) on a Minispec+ analyzer (Bruker, Billerica, MA, USA) on conscious mice at the age of 7 weeks (while on CD) and 20 weeks (after 11 weeks of HFD).

#### 2.8.2. Glucose tolerance test

At the age of 21 weeks, an oral glucose tolerance test (OGTT) was performed after 4 hours’ fasting. After measurement of the basal glucose level using a drop of blood collected from the tail (time 0), mice were administered per os with a solution of 20% glucose in sterile saline at a dose of 1 g glucose/kg body weight. Blood was collected for glucose determination from the tail 20, 40, 60, and 90 min after injection of the glucose solution. The incremental area of the glucose curve was then calculated as a measure of glucose tolerance.

#### 2.8.3. Insulin tolerance test

At the age of 22 weeks, insulin sensitivity was evaluated via an intraperitoneal insulin tolerance test (ipITT) on animals that were fasted for 2 hours. After measurement of the basal glucose level, using a drop of blood collected from the tail (time 0), mice were injected intraperitoneally with a solution of insulin (Umuline rapide [Lilly, Indianapolis, IN, USA]) at a dose of 1 U/kg body weight. Blood was collected for glucose determination from the tail 15, 30, 45, 60, and 90 min after injection of insulin.

#### 2.8.4. Blood collection and analysis

Lipids (triglycerides, free fatty acids, glycerol, total cholesterol, HDL cholesterol and LDL cholesterol) and glucose were measured in plasma from blood collected by retro-orbital puncture under isoflurane anaesthesia at the age of 9 and 25 weeks. At 25 weeks of age, a complete blood chemistry analysis and a complete blood cell count were performed, including electrolytes and ions (sodium, potassium, chloride, calcium, phosphorus, magnesium), metabolites (urea, creatinine, total proteins, albumin, total bilirubin, bile acids) and enzymes (alanine aminotransferase [ALAT], lactate dehydrogenase, alkaline phosphatase alpha-amylase). These parameters were determined on an OLYMPUS AU-400 automated laboratory work station (Beckman Coulter, Brea, CA, USA), with kits and controls supplied by Beckmann Coulter (Beckman Coulter, Lismeehan, Ireland). Free fatty acid was measured on the AU-400 using a kit from Wako (Wako Chemical Inc., Richmond, VA, USA). Glycerol was measured using a kit from Randox (Randox Laboratories, Crumlin, UK).

### 2.9. Effects of acute and subchronic pharmacological BACE inhibition

The effects of acute and subchronic BACE inhibition in WT mice were investigated by single- dose MBi-7 and 3-week oral administration of MBi-3, respectively. Further details are given in the ESM.

### 2.10. Statistical analyses

Unless otherwise stated, errors for normalized data are expressed as SD due to propagation, and errors for non-normalized data are expressed as SEM. All data were tested for normality using the Anderson-Darling test. Parameters that demonstrated non-normal data in any one of the three experiments were excluded from parametric statistical analysis and were analyzed by non-parametric Mann–Whitney U testing [30]. Parametric testing was performed using Student’s t-test. Outliers were excluded if their values exceeded two SDs away from the mean. All p-values reflect two-tailed testing, with the method of analysis provided in the appropriate legends.

## 3. Results

### 3.1. KO mouse-line characterisation

Analysis of BACE proteins, TMEM27 processing and pancreatic islet profiles in the *Bace1* and/or *Bace2* KO mice are given in the ESM (Supplementary Results and Fig. S3–S5).

### 3.2. Mice carrying *Bace1* mutations do not respond to HFD challenge

A HFD had only a small impact on female mice. Therefore, we focused our analyses on males, in which a HFD response was evident in WT animals; results for female mice are provided in the ESM (Supplementary Results and Fig. S6–S8).

At 5 weeks of age, mice lacking BACE1, i.e. *Bace1* KO and *Bace1/2* dKO, weighed less than WT counterparts (Fig. 1a). On CD, these male mice demonstrated a statistically significant increase of body weight until week 9, when the mice were fed a HFD. After 2 weeks on a HFD, two *Bace1* KO male mice were found dead in their cages, with no apparent link to genotype.

**Fig. 1.**
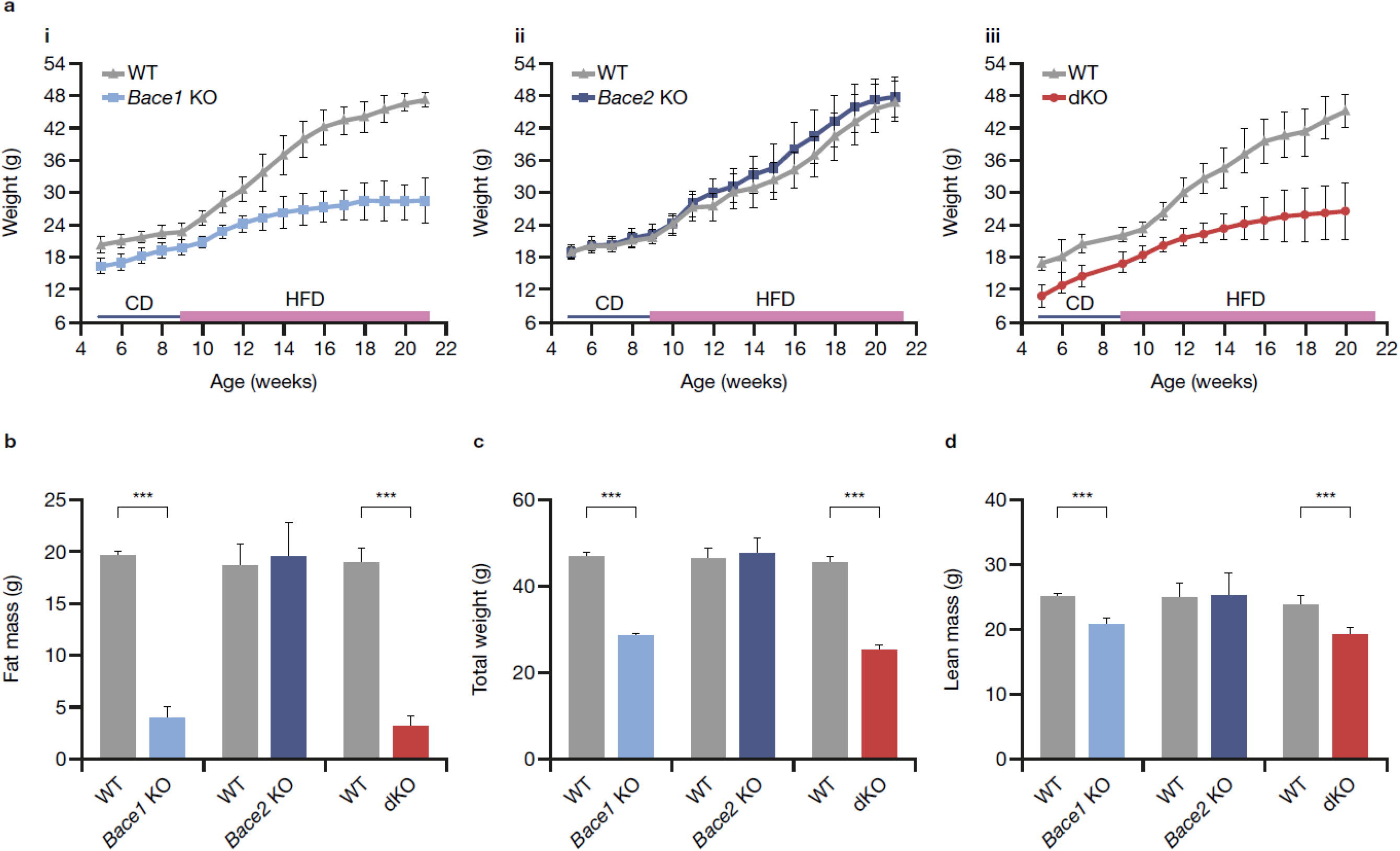
Differential response of *Bace* mutants to HFD challenge for growth and body composition. (**a**) Weight-evolution curves of male mice for experiments involving i) *Bace1* KO, ii) *Bace2* KO and iii) dKO mutants. (**b**) Fat mass as determined by body composition analysis in male mice. All data are presented as averages ± SD. Statistical analyses performed using Mann–Whitney U test for genotype effect. (**c**) Weights at the time of post-HFD–challenge body composition analysis in male mice. (**d**) Lean mass as determined by body composition analysis in male mice. Statistical analyses performed using Mann–Whitney U test for genotype effect. *\*p* < 0.05; *\*\*\*p* < 0.001. *Bace*, beta-site amyloid precursor protein-cleaving enzyme; CD, chow diet; dKO, double *Bace1*/*Bace2* knockout; HFD, high-fat high-cholesterol diet; KO, knockout; WT, wild-type

These individuals were excluded from all HFD analyses. Upon HFD challenge, WT males in each of the three cohorts showed consistent weight gain of approximately 100% throughout the course of the studies. *Bace2* KO males showed weight gain similar to WT animals (Fig. 1a). In contrast, the *Bace1* KO and *Bace1/2* dKO males showed only ∼25% weight increase, which was highly statistically significantly different to WT. The age dependence of weight gain for the *Bace1* KO and *Bace1/2* dKO males was similar on CD and HFD (Fig. 1a).

### 3.3. *Bace1* KO mice show defects in fat deposition

Before HFD challenge, both the *Bace1* KO and *Bace1/2* dKO mice showed highly statistically significant differences from WT in lean mass, consistent with their reduced weights (ESM; Fig. S6b); however, there was no statistically significant difference in fat composition for any of the mutant mice compared with WT by qNMR (ESM; Fig; S6b).

After HFD challenge, *Bace2* KO male mice acquired fat deposits similar to their WT counterparts (Fig. 1b), resulting in no statistically significant difference in total weight between groups (Fig. 1c). In contrast, *Bace1* KO and *Bace1/2* dKO males had dramatically less fat mass accumulation than WT (Fig. 1b), which accounted for the majority of the difference in total weight between these groups, although there was also a continued reduction in lean mass (Fig. 1d).

### 3.4. Loss of BACE1, but not BACE2, induces improved glucose tolerance

*Bace1* KO and *Bace1/2* dKO male mice had highly statistically significantly lower fasted blood glucose levels and reduced amplitude of blood glucose increase compared with WT mice (*p* < 0.001) (Fig. 2a), whereas *Bace2* KO and WT males had similar blood glucose kinetics. When the effect of administering a glucose bolus was measured as a percentage change from fasted blood glucose levels (Fig. 2b), the effects on *Bace1* KO and dKO mice were less pronounced but the *Bace1* KO and dKO male mice showed statistically significant improved glucose tolerance at later time-points compared with WT mice (*p* < 0.05), while no effect of *Bace2* KO was apparent.

**Fig. 2.**
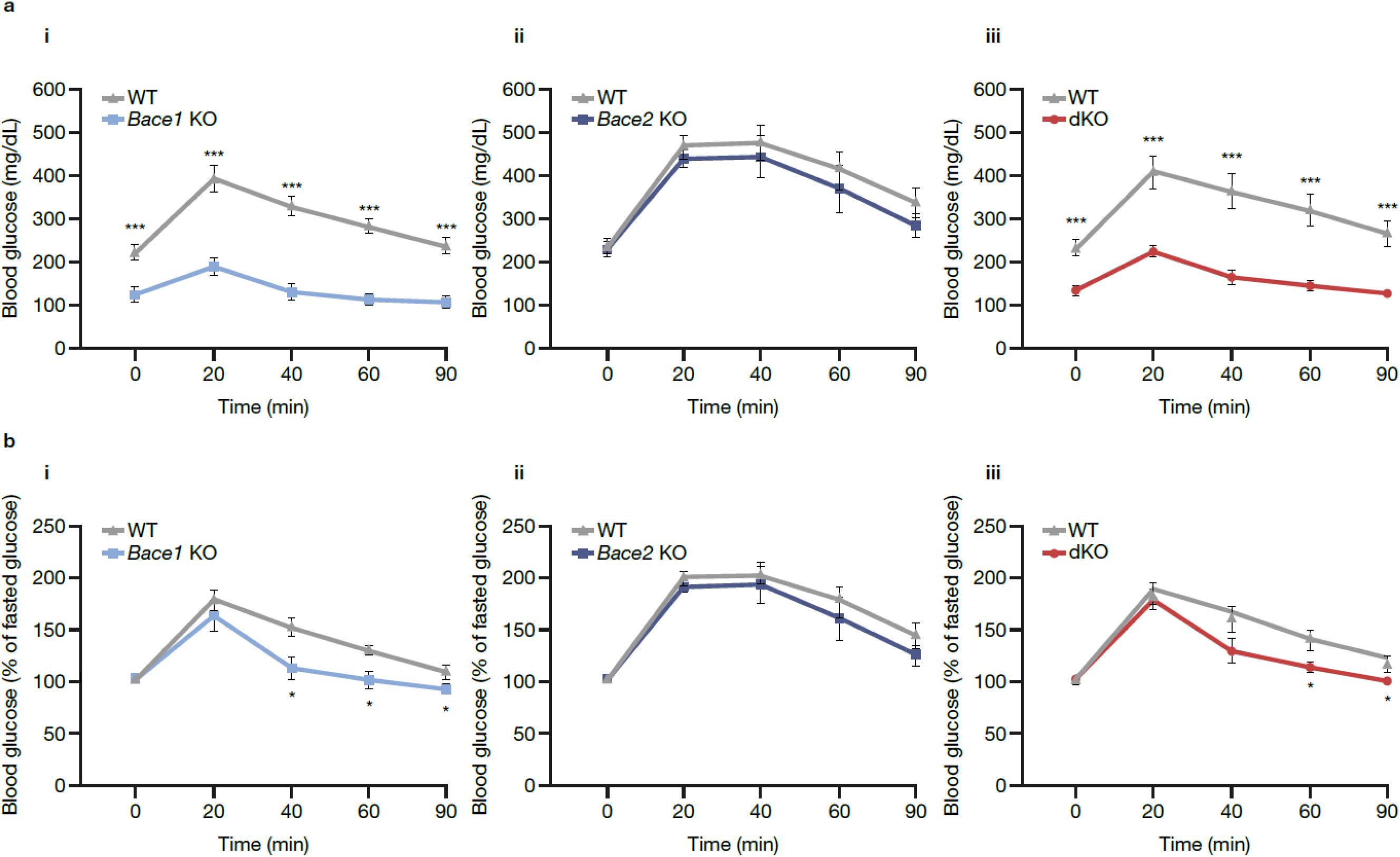
Glucose tolerance testing of *Bace* mutant mice after HFD challenge. (**a**) Raw blood glucose measurement from fasting (t = 0) and time after glucose bolus (t = 20–90 min) for males of each of the cohorts examined: i) *Bace1* KO, ii) *Bace2* KO and iii) dKO mutants. (**b**) Blood glucose measurements as a percentage of fasted blood glucose for individual mice: i) *Bace1* KO, ii) *Bace2* KO and iii) dKO mutants. All data are presented as averages ± SEM. Statistical analysis performed by Student’s t-test for genotype effect. *\*p* < 0.05; *\*\*\*p* < 0.001. *Bace*, beta-site amyloid precursor protein-cleaving enzyme; dKO, double *Bace1*/*Bace2* knockout; HFD, high-fat high-cholesterol diet; WT, wild-type

### 3.5. Loss of BACE1, but not BACE2, improves insulin sensitivity

ipITT tests on HFD-challenged male mice demonstrated greater variability between WT controls in the three cohorts (Fig. 3). However, both *Bace1* KO and dKO showed consistently depressed blood glucose levels, with improved insulin sensitivity with respect to fasted blood glucose levels (Fig. 3a), even when values were corrected for percentage of fasted glucose (Fig. 3b). In contrast, *Bace2* KO mice showed no statistically significant difference relative to WT.

**Fig. 3.**
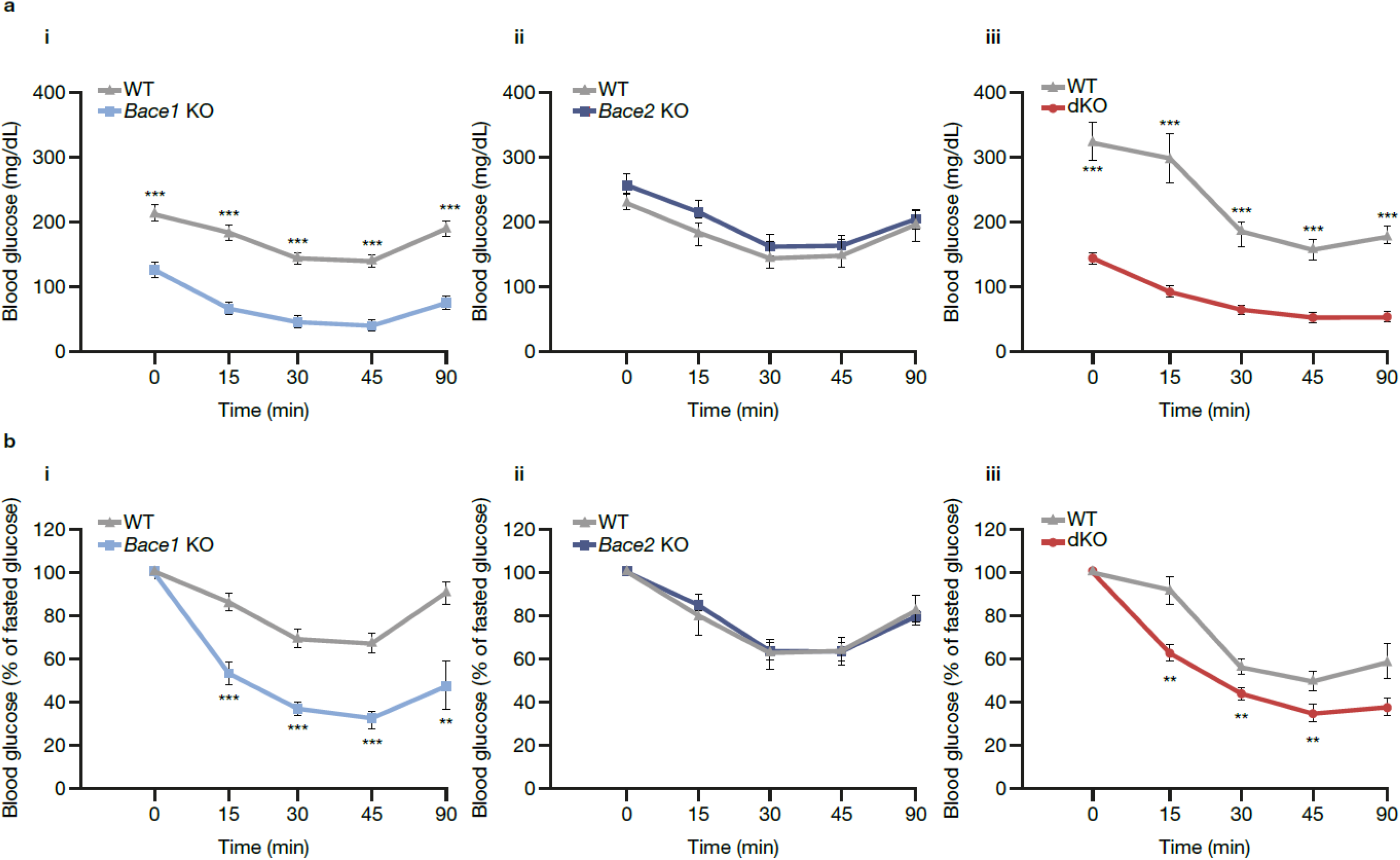
Insulin sensitivity testing of *Bace* mutant mice after HFD challenge. (**a**) Raw blood glucose measurement from fasting (t = 0) and time after intraperitoneal insulin injection (t = 15–90 min) for males of each of the cohorts examined: i) *Bace1* KO, ii) *Bace2* KO and iii) dKO mutants. (**b**) Blood glucose measurements as a percentage of fasted blood glucose for individual mice: i) *Bace1* KO, ii) *Bace2* KO and iii) dKO mutants. All data are presented as averages ± SEM. Statistical analysis performed by Student’s t-test for genotype effect. *\*\*p* < 0.01; *\*\*\*p* < 0.001. *Bace*, beta-site amyloid precursor protein-cleaving enzyme; dKO, double *Bace1*/*Bace2* knockout; WT, wild-type

### 3.6. Loss of BACE1, but not BACE2, reduces diet-induced hypercholesterolaemia

Cholesterol parameters for *Bace1* KO and dKO mice following HFD were highly statistically significant different to those of WT mice (Fig. 4). Compared with pre-HFD, 2-fold to 3-fold increases in total, HDL and LDL cholesterol were observed in WT and *Bace2* KO male mice (WT: total cholesterol, 2.37–2.69 vs. 5.70–6.26 mmol/L; HDL cholesterol, 1.56–1.87 vs. 3.41–4.12 mmol/L; LDL cholesterol, 0.37–0.44 vs. 1.16–1.23 mmol/L; *Bace2* KO: total cholesterol, 2.51 vs.5.75 mmol/L; HDL cholesterol, 1.67 vs.3.81 mmol/L; LDL cholesterol, 0.45 vs.1.13 mmol/L). In contrast, *Bace1* KO and dKO mice only demonstrated an approximate 0.5-fold increase in these parameters (*Bace1* KO: total cholesterol, 2.16 vs.3.05 mmol/L; HDL cholesterol, 1.37 vs.2.12 mmol/L; LDL cholesterol; 0.36 vs.0.54 mmol/L; dKO: total cholesterol, 2.43 vs.3.18 mmol/L; HDL cholesterol, 1.67 vs.1.92 mmol/L; LDL cholesterol, 0.43 vs.0.67 mmol/L).

**Fig. 4.**
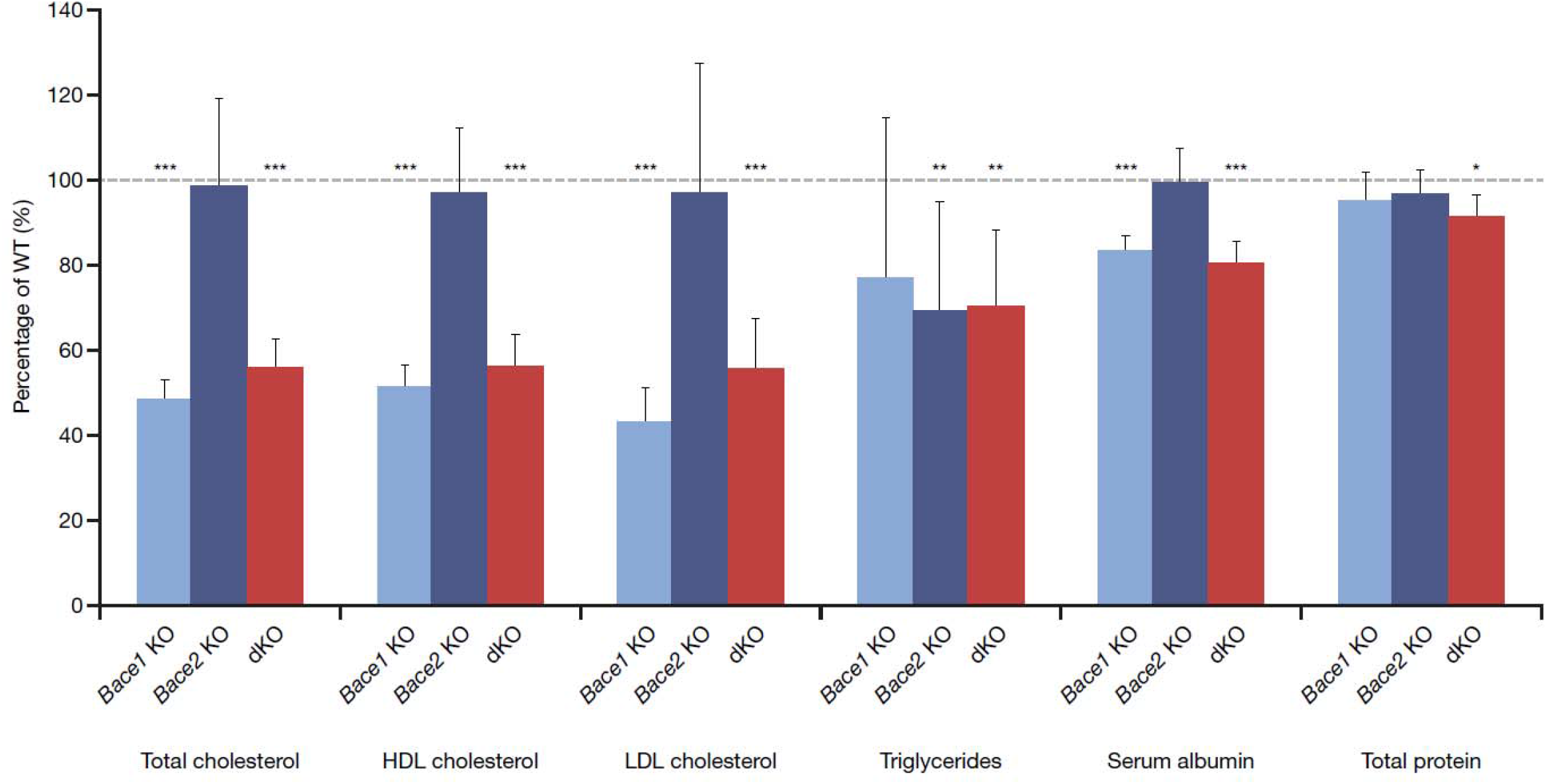
Effects of *Bace* mutants on blood chemistry following HFD challenge. Blood chemistry parameters for the male mice of the three *Bace* cohorts examined. Mutant values are normalized to WTs analyzed concurrently. Data are expressed as averages ± SD. Statistical analyses performed using Mann–Whitney U test for genotype effect. *\*p* < 0.05; *\*\*p* < 0.01; *\*\*\*p* < 0.001. *Bace*, beta-site amyloid precursor protein-cleaving enzyme; dKO, double *Bace1*/*Bace2* knockout; WT, wild-type

The dKO male mice did not show the same degree of triglyceride increase after HFD challenge as WT mice. Interestingly, a similar effect was seen for both *Bace2* KO and *Bace1* KO, although the effect in *Bace1* KO mice was not statistically significantly different to WT (Fig. 4). *Bace1* KO and dKO mice also demonstrated similarly low serum albumin – an indicator of liver dysfunction and/or deficient protein uptake – after HFD challenge (Fig. 4). No effect was observed on ALAT activity (data not shown).

### 3.7. Reduced HFD-induced liver fat deposition in the absence of BACE1

Whereas no statistically significant differences in weight of the kidneys, spleen or heart at necropsy were noted in male mice (data not shown), a highly statistically significant difference in liver weight was noted for *Bace1* KO and dKO mice vs. WT, but not for *Bace2* KO male mice (Fig. 5a). Fat deposits were found in the livers of WT and *Bace2* KO male mice after HFD challenge, but not in the livers of *Bace1* KO or dKO males (Fig. 5b).

**Fig. 5.**
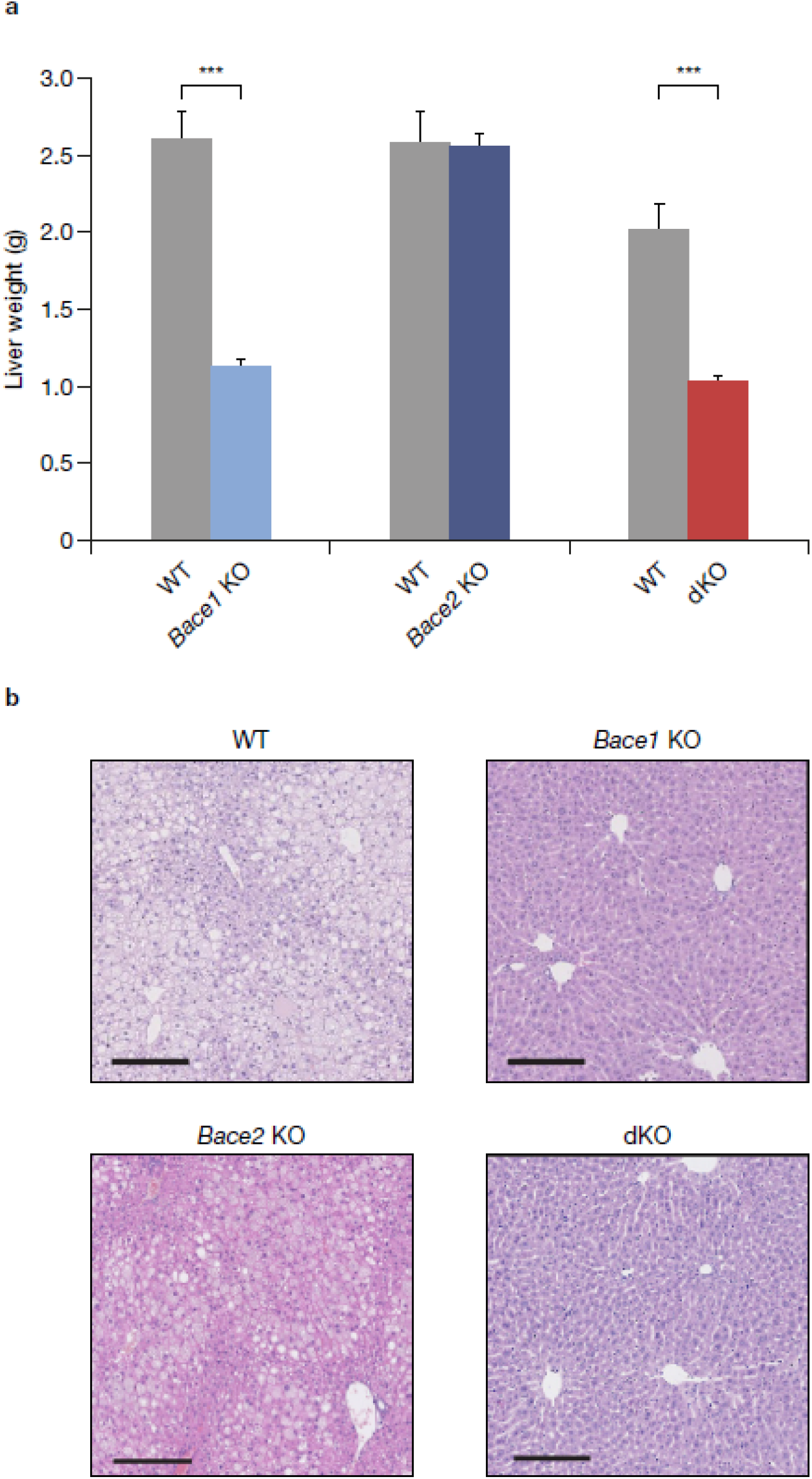
BACE1, but not BACE2, is important for liver fat deposition during HFD challenge. (**a**) Liver weights at necropsy following HFD challenge in male mice. All data are presented as averages ± SEM. Statistical analyses performed using Mann–Whitney U test for genotype effect. *\*\*\*p* < 0.001. (**b**) Representative images of livers visualized by haematoxylin and eosin staining for each of the genotypes analyzed. Size bar = 200 μm. *Bace*, beta-site amyloid precursor protein-cleaving enzyme; dKO, double *Bace1/Bace2* knockout; HFD, high-fat high-cholesterol diet; WT, wild-type

### 3.8. Effect of acute and subchronic BACE inhibition on lean mice

Oral administration of MBi-7 (ESM; Fig. S2), which has ∼3-fold higher affinity for BACE2 (K_i_: 17 nM) than for BACE1 (K_i_: 57 nM), to lean mice resulted in a significant increase in plasma glucagon-like peptide 1 (GLP-1) concentrations following glucose challenge (Fig. 6a). Similarly, insulin plasma concentrations increased significantly in comparison with the vehicle-treated group post-glucose challenge (Fig. 6b). Subchronic inhibition by MBi-3 resulted in no changes in body-weight growth or basal blood glucose relative to age-matched HFD-fed controls.

**Fig. 6.**
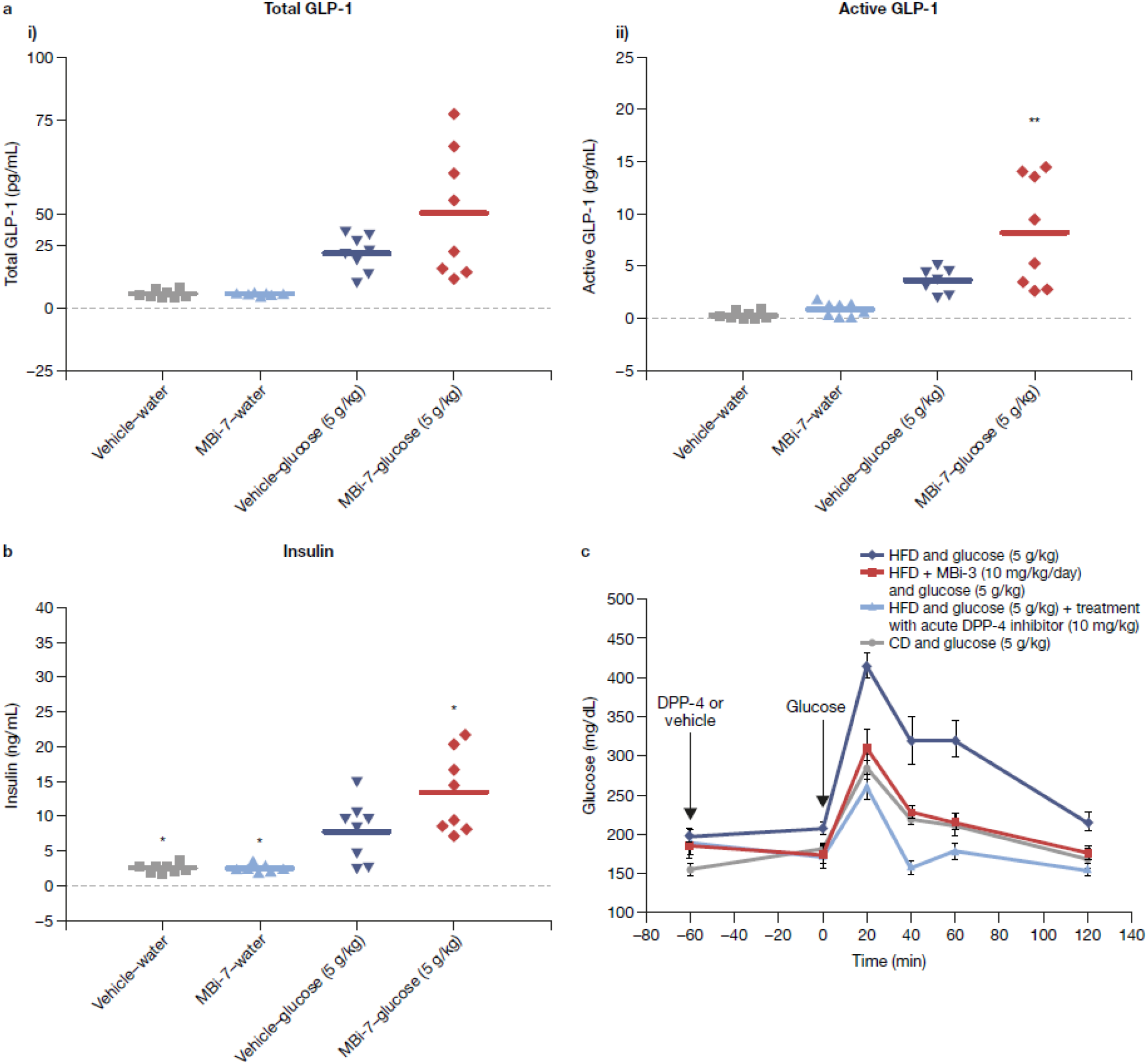
Acute effect of MBi-7 on plasma GLP-1 and insulin concentrations and effect of subchronic treatment of MBi-3 on blood glucose excursion in OGTT in 3-week HFD mice. C57BL/6N mice were fasted for 7 hours prior to treatment with vehicle or MBi-7 at 30 mg/kg. At 2 hours post-treatment, mice in both groups were challenged with either water or glucose (5 g/kg body weight). After 10 min, i) total and ii) active GLP-1 (**a**) and insulin (**b**) concentrations were measured. Statistical analysis performed using ANOVA. (**c**) C57BL/6N mice were treated for 3 weeks either on a HFD, HFD + MBi-3 mixed in food, HFD + acute treatment with DPP-4 inhibitor desfluoro-sitagliptin, or on regular CD, and blood glucose levels measured after glucose challenge (5 g/kg glucose). ANOVA analysis: *\*p* < 0.05; *\*\*p* < 0.01; *\*\*\*p* < 0.001 vs. vehicle. ANOVA, analysis of variance; CD, chow diet; DPP-4, dipeptidyl peptidase 4; GLP-1, glucagon-like peptide 1; HFD, high-fat high-cholesterol diet; OGTT, oral glucose tolerance test

Evaluation after 3 weeks on diet (19 weeks of age) revealed a modest reduction in blood glucose concentrations at 1-hour post-glucose administration in the MBi-3 and dipeptidyl peptidase-4 (DPP-4) inhibitor (positive control) groups, as well as improved glucose tolerance post-glucose challenge in an OGTT (Fig. 6c). MBi-3 treatment corrected glucose excursion to the response observed in mice maintained on a lean diet.

## 4. Discussion

In the present study, we compared the metabolic effects of the loss of BACE1, BACE2 or both in KO mice after a 16-week treatment on a HFD. We also investigated changes in blood parameters such as cholesterol, and the histological consequences of a HFD on mutant and WT mice and in mice treated with BACE inhibitors.

We found that *Bace1* KO and dKO mice – but not *Bace2* KO mice – had significantly lower body weight but no difference in fat composition compared with WT controls at 5 weeks of age, suggesting a growth delay consistent with previous KO studies [21]. The weight gain observed during HFD challenge in *Bace2* KO and WT males but not *Bace1* KO and dKO males indicates a lack of effect of dietary constituents on weight gain in *Bace1* KO and dKO animals, suggesting that HFD-induced weight gain is dependent on BACE1 functioning. Following administration of a HFD, a regimen known to sensitize mice to metabolic disorders [31], *Bace1* KO and dKO mice had also dramatically less fat mass than *Bace2* KO or WT mice, suggesting that diet-induced deposition of fat requires BACE1, but not BACE2, expression and function. A similar effect was observed in the fat deposits in the liver. Similar blood glucose kinetics in *Bace2* KO and WT male mice were observed throughout, whereas *Bace1* KO and dKO male mice showed lower fasted blood glucose and a lower amplitude of blood glucose increase. The ipITT revealed highly significantly depressed blood glucose levels, with improved insulin sensitivity in HFD-treated *Bace1* KO and dKO males compared with *Bace2* KO and WT males. Based on the findings of the ipITT and OGTT, we conclude that BACE1 is important in glucose homeostasis after diet-induced challenge and that BACE2 is dispensable in this function.

*Bace1* KO and dKO males had lower levels of total and HDL cholesterol than *Bace2* KO and WT males on a CD, and demonstrated smaller increases in total, HDL and LDL cholesterol after HFD challenge, suggesting that BACE1 is necessary for the full impact of diet-induced hypercholesterolaemia. Increased triglycerides after HFD challenge required BACE2; however, this parameter does not appear to impact the other physiological effects of HFD challenge that are not changed in *Bace2* KO male mice. Although both *Bace1* KO and dKO males showed similarly low serum albumin after HFD challenge, suggesting that the physiological impacts of the dKO are mostly attributable to the *Bace1* deficiency, the lack of an effect on ALAT may reflect a difference in nutritional uptake rather than liver damage, consistent with the absence of severe HFD-induced weight gain. Liver damage previously reported for a subset of dual BACE inhibitors has been shown to be structure- rather than mechanism-dependent [32], and no hepatotoxicity was reported in a previous preclinical study of verubecestat [9].

Hypercholesterolaemia can impair the cholinergic system and lead to memory deficits in rodents [33], and may contribute to the onset of AD-like dementia, increased BACE1 expression, and acceleration of Aβ-induced cognitive deficits [34, 35]. Thus, it is notable that our data support a role of BACE1 function in the onset of hypercholesterolaemia.

Our data corroborate earlier findings that loss of BACE1 function results in decreased body weight gain, increased glucose tolerance and enhanced insulin sensitivity [21]. Interestingly, the underweight phenotype of the *Bace1* KO mice was greatly reduced and completely absent in post-natal forebrain excitatory neurons and adult whole-body conditionally *Bace1* KO mice, respectively [36]. Decreased BACE1 activity also normalizes hypothalamic inflammation, lowers protein tyrosine phosphatase 1B (PTP1B) and suppressor of cytokine signalling 3 (SOCS3) – both negative regulators of leptin – and restores hypothalamic leptin sensitivity in obese, but not lean, mice [37]. Conversely, mice overexpressing human *BACE1* in the forebrain develop systemic diabetes, supporting a role for BACE1 in metabolism regulation through a centrally mediated process [22].

The BACE2 protein tissue-expression profile described here is consistent with the tissue-expression profile of *BACE2* RNA as defined from multiple sources (e.g. https://www.proteinatlas.org/ENSG00000182240-BACE2/tissue). Furthermore, our data confirm that BACE2 is indeed the major sheddase for TMEM27, with MBi-2 inhibiting formation of secreted TMEM27 from full-length TMEM27 in a dose-dependent manner *in vitro* and elevated full-length TMEM27 levels in the *Bace2* KO mice compared with WT mice, similar to a previous report of *ad-libitum*–fed *Bace2*-Δ6/Δ6-deletion mutant mice, which expressed functionally inactive BACE2 [20]. *Bace2*-Δ6/Δ6-deletion mice also exhibited increased beta-cell mass due to increased beta-cell proliferation [20], while *Bace2* KO mice had increased beta-cell numbers and reduced TMEM27 processing associated with improved glucose tolerance [38]. In human islet amyloid polypeptide-transgenic (hIAPP-Tg) mice, *BACE2* KO promoted beta-cell survival and function and reverted hIAPP-Tg–associated glucose intolerance and decreased insulin secretion [23], which is in line with the glucose tolerance data presented here. No changes in islet area or proliferation were evident in the current *Bace2* KO mice compared with WT mice, in contrast to *Bace2*-Δ6/Δ6-deletion mutant mice, which also had decreased blood glucose levels and improved glucose tolerance on a regular CD compared with WT mice [20], and may have further contributed to the lack of effects of BACE2 on OGTT and ipITT endpoints. The differences in beta-cell density between the two *Bace2* mouse lines may be due to subtle genetic differences in background strains, animal husbandry conditions such as type of chow or mouse microbiome, or due to the fact that the *Bace2* KO line described here is devoid of BACE2 protein expression, whereas the *Bace2*-Δ6/Δ6 line (Esterházy et al. 2011) retains expression of a truncated, inactive, soluble BACE2 protein that may have unexpected impacts on phenotypes, perhaps via aberrant dimerisation with BACE1 and suppression of its function [39].Our finding that loss of BACE1 protects against HFD-induced obesity and diabetes suggests that chronic pharmacological inhibition with BACE inhibitors may protect against HFD-induced metabolic challenges. Indeed, acute BACE inhibition in lean mice resulted in a significant increase in plasma GLP-1 – an incretin hormone that enhances insulin and suppresses glucagon secretion [40] – and insulin concentrations vs. controls after glucose challenge.

However, subchronic BACE inhibition in HFD-fed mice resulted in no changes in body weight or basal blood glucose, and a modest reduction in blood glucose concentrations and improved glucose tolerance, while acute treatment with desfluoro-sitagliptin resulted in a robust improvement of glucose tolerance, as expected. Pharmacological BACE inhibition has been shown to result in reduced body weight, improved glucose homeostasis and decreased plasma leptin levels in diet-induced obese mice after a 20-week HFD [37]; therefore, longer treatment may result in a more profound treatment effect. However, it is unlikely that BACE inhibition is a viable therapeutic strategy to treat type 2 diabetes, as the effect size appears to be relatively small in comparison with alternative approaches such as GLP-1 agonists and DPP-4 inhibitors, which are routinely used to treat type 2 diabetes [40–42].

A promising neuroprotective effect of GLP-1 has been observed in animal models of AD [43, 44], which is notable given the link between insulin resistance and cognitive deficits seen in AD [45, 46]. Brain levels of insulin and insulin receptor were lower in AD brains and animal models of AD [47]. Therefore, therapeutic approaches, such as decreased production of toxic Aβ oligomers via BACE inhibition, may reduce impaired insulin signalling in the brain [48]. The dual BACE inhibitor verubecestat (MK-8931) has been evaluated in Phase 3 clinical trials for treatment of patients with mild-to-moderate and prodromal AD [15, 16]. Notably, long-term (78 and 104 weeks) treatment with verubecestat resulted in a modest but significant weight loss vs. a weight increase in the placebo group; however, the relative contribution of changes in fat vs. lean mass to body weight changes was not measured. [15, 16].

## 5. Conclusions

Our data indicate that lack of BACE1 – but not BACE2 – function contributes mainly to the metabolic phenotypic changes observed in *Bace1/2* dKO mice, suggesting that inhibition of BACE1 has the greater role (vs. BACE2) in any potential improvements in metabolic homeostasis that occur as a result of loss of BACE activity.

## ABBREVIATIONS

β: amyloid beta
AD: Alzheimer’s disease
ALAT: alanine aminotransferase
ANOVA: analysis of covariance
APP: amyloid precursor protein
BACE: beta-site amyloid precursor protein-cleaving enzyme
CD: chow diet
CSF: cerebrospinal fluid
DAPI: 4 ,6-diamidino-2-phenylindole
dKO: double *Bace1*/*Bace2* knockout
DMEM: Dulbecco’s modified Eagle’s medium
DMSO: dimethyl sulfoxide
DNA: deoxyribonucleic acid
DPP-4: dipeptidyl peptidase-4
ES: embryonic stem
ESM: electronic supplementary material
FBS: fetal bovine serum
GLP-1: glucagon-like peptide 1
HET: heterogeneous
HFD: high-fat high-cholesterol diet
hIAPP-Tg: human islet amyloid polypeptide-transgenic
ipITT: intraperitoneal insulin tolerance test
KO: knockout
nd: not detected
OGTT: oral glucose tolerance test
PTP1B: protein tyrosine phosphatase 1B
qNMR: quantitative nuclear magnetic resonance
TMEM27: transmembrane protein 27
WT: wild-type

## Acknowledgments

Medical writing assistance, under the direction of the authors, was provided by Kirsty Muirhead, PhD, of CMC AFFINITY, McCann Health Medical Communications, in accordance with Good Publication Practice (GPP3) guidelines. This assistance was funded by Merck Sharp & Dohme Corp., a subsidiary of Merck & Co., Inc., Kenilworth, NJ, USA.

## Data availability statement

Merck Sharp & Dohme Corp., a subsidiary of Merck & Co., Inc., Kenilworth, NJ, USA’s data sharing policy, including restrictions, is available at http://engagezone.msd.com/ds_documentation.php. Requests for access to study data can be submitted through the EngageZone site or via email to.

## Contribution statements

TWR: contributed to conception of the research, design of mouse lines, *in vivo* experimental designs, data analysis, figure preparation, and writing of the manuscript.

LAH: contributed to conception of the research, *in vivo* experimental designs, data analysis, and writing of the manuscript.

PTR: contributed to data analysis, figure preparation, and writing of the manuscript.

M-FC: contributed to *in vivo* experimental designs, data acquisition and analysis, figure preparation, and writing of the manuscript.

KJB: contributed to design of *in vivo* experiments, data acquisition and analysis, figure preparation, and writing of the manuscript.

BP-D: contributed to *in vivo* experimental designs, data acquisition and analysis, figure preparation, and writing of the manuscript.

TS: contributed to *in vivo* experimental designs, data acquisition and analysis, figure preparation, and writing of the manuscript.

HJ: contributed to *in vivo* experimental designs, data acquisition and analysis, figure preparation, and writing of the manuscript.

RT: contributed to experimental designs, data acquisition, data analysis, figure preparation, and writing of the manuscript.

JDS: Contributed to the design and synthesis of BACE inhibitors and writing of the manuscript.

JNC: Contributed to the design and synthesis of BACE inhibitors and writing of the manuscript.

EMP: Contributed to conception of the research and writing of the manuscript.

MEK: Contributed to conception of the research, *in vitro* experimental designs, data collection analysis, and writing of the manuscript.

All authors approved the manuscript for submission.

## Electronic supplementary material

### 1. Supplementary Methods

#### 1.1. Generation of *Bace1* KO, *Bace2* KO and *Bace1/2* dKO mice

In the *Bace1* knockout (KO) mice, exon 1 was replaced by homologous recombination in embryonic stem (ES) cells using a targeting vector carrying the flippase recognition target (FRT)-site–flanked neomycin-resistance gene cassette, resulting in a full null allele (Fig. S1a). Similarly, a targeting vector introduced a deletion in exon 1 in the *Bace2* KO mice (Fig. S1b). Correctly targeted ES-cell clones were microinjected into blastocysts and the resulting chimeric mice further bred with C57BL/6N mice for germline transmission of the targeted alleles. *Bace1* and *Bace2* KO mice were established by crossing heterozygous mice, and *Bace1/2* double-KO (dKO) mice by crossing *Bace1* and *Bace2* single-KO mice.

#### 1.2. Preparation of samples for determination of BACE1 and BACE2 expression in *Bace2* KO mouse tissue

Tissues from wild-type (WT) and *Bace2* KO mice were stored at –80°C prior to preparing membrane protein fractions. Tissues were homogenized in 1 mL of 20 mM HEPES, pH 7.4, 2 mM EDTA, 2 mM EGTA, 50 mM KCl, 250 mM sucrose, protease inhibitor cocktail (Sigma-Aldrich, St. Louis, MO, USA) (buffer A) using a polytron (20,000 rpm, 30 seconds), and placed on ice. Insoluble material was pelleted (15,000 × *g*, 30 min), resuspended in buffer A containing 1% Triton X-100 and agitated for 1 hour at 4°C. Supernatants were collected after pelleting insoluble material (100,000 × *g*, 30 min), and protein concentrations determined by a detergent-compatible Assay (BioRad, Hercules, CA, USA).

#### 1.3. Pharmacological BACE inhibition

##### 1.3.1. Acute effect of the BACE inhibitor MBi-7 on plasma insulin and glucagon-like peptide 1 (GLP-1) concentrations

11-week-old male WT mice were housed 8 per cage with *ad libitum* access to food and water. On the morning of the study, mice were randomized (*n* = 8 per group) and fasted for 7 hours prior to treatment with vehicle (0.5% methylcellulose [5 mL/kg]) or MBi-7 (30 mg/kg). At 2 hours post-treatment, mice in both study groups were orally challenged with either water (10 mL/kg) or glucose (5 g/kg body weight as a 5% solution). At 10 min post-challenge, mice were euthanized by CO_2_ asphyxiation and plasma was harvested from cardiac puncture bleeds and collected into a protease inhibitor cocktail (Sigma-Aldrich). Total and active GLP-1 plasma concentrations were measured using kits K110FCC-2 and K110HZC-2 (Meso Scale Diagnostic, Rockville, MD, USA), respectively. Insulin was measured using enzyme-linked immunosorbent assay kit 10- 1250-10 (Mercodia, Winston Salem, NC, USA).

##### 1.3.2. Effect of subchronic treatment of the BACE inhibitor MBi-3 on body weight, ambient blood glucose and blood glucose excursion in an oral glucose tolerance test (OGTT) in 3-week high-fat high-cholesterol diet (HFD)-fed mice

11-week-old male WT mice (*n* = 5/cage) were randomized by weight and distributed into four treatment groups (*n* = 10 per group). A HFD control group and the acute dipeptidyl peptidase-4 (DPP-4) treatment group had *ad libitum* access to water and a HFD (M12492MI, 60% kcal fat; Research Diets, New Brunswick, NJ, USA). Stable BACE inhibition (central nervous system Aβ effective dose in 75% of the population [ED_75_]) was achieved via MBi-3 ad-mixed into this diet and fed to lean mice for 4 weeks of treatment to provide an exposure of 10 mg/kg/day based on daily food intake and body weight. A lean control group was maintained on Teklad 7012 diet (Envigo, Huntingdon, UK). Body weight and ambient blood glucose (tail nick/ultra one-touch glucometer, LifeScan, Wayne, PA, USA) were monitored weekly during the 3-week treatment period prior to the OGTT. On the morning of the OGTT, mice were fasted for 6 hours prior to glucose challenge. At 1 hour prior to glucose challenge (t = –60 min), baseline blood glucose was determined (tail nick/glucometer) and the lean control, MBi-3 treatment group and HFD control mice were administered an oral dose of vehicle (0.5% methylcellulose at 5 mL/kg). The acute DPP-4 inhibitor group received a maximally efficacious acute oral dose of the DPP-4 inhibitor desfluoro-sitagliptin. At t = 0, blood glucose was determined, and then all groups were orally administered glucose (5 g/kg in water at 10 mL/kg). Blood glucose was monitored at 20, 40, 60, and 120 min post-glucose challenge to profile the effects of treatments on blood glucose excursion.

### 2. Supplementary Results

#### 2.1. *Bace1 and Bace2* KO mice lack BACE proteins

Brain lysates from *Bace1* heterozygous mice displayed reduced BACE1 protein relative to WT, while BACE1 protein was undetectable in the brain lysates of *Bace1* KO mice (Fig. S1a, right). A summary of phenotypic profiling of this *Bace1* KO line will be published separately. To verify the loss of BACE2 protein expression in the *Bace2* KO mice, we identified commercially available antibodies specific for BACE2 vs. BACE1 by western blotting of whole-cell extracts from HEK293 cells transiently overexpressing mouse BACE1 and mouse BACE2 (data not shown). Expression of a single ∼60 kDa protein in WT intestinal tissue but not WT brain or *Bace2* KO intestinal or brain tissue further confirmed the antibody specificity for BACE2 (Fig. S1b, right). Significant and specific BACE2 expression was observed in the gastrointestinal tract, trachea, ovary, bladder and lung of WT but not *Bace2* KO mice (Fig. S3). The same tissues showed clear expression of BACE1 only in brain, indicating BACE1 protein expression was not upregulated in *Bace2* KO mice (Fig. S3). No BACE2 band was detected in pancreatic tissue despite known *Bace2* RNA and protein expression in pancreatic beta-cells [18, 19].

However, the more sensitive technique of immunofluorescence staining clearly detected BACE2 protein in fixed islets from WT but not *Bace2* KO mice (Fig. S4b). Additionally, protein bands at the approximate molecular weight of BACE2 were present in both WT and KO extracts from liver. The absence of the 60 kDa band in multiple tissues from *Bace2* KO animals confirmed antibody specificity for BACE2 and the absence of BACE2 protein in the *Bace2* KO mouse (Fig. S3 and S4c).

BACE2-dependent processing of TMEM27 in cells and loss of BACE2 processing of TMEM27 in acutely isolated primary cultured islets from *Bace2* KO mice was confirmed. Western blot analysis of HEK293 cells transfected with mouse *Tmem27* detected a TMEM27 immunoreactive protein species of ∼40–42 kDa, consistent with full-length TMEM27, in whole-cell lysates and a weakly reactive ∼25–28 kDa band, likely arising from the BACE2-mediated shedding of a soluble N-terminal fragment of TMEM27, in the cell culture supernatant (Fig. S4c, left). Co-transfection of *Tmem27* with human *BACE1* did not increase secreted TMEM27 protein, while co-transfection with human *BACE2* resulted in the loss of detectable full-length TMEM27 and the appearance of secreted TMEM27 protein in the cell-culture supernatant (Fig. S4c, left).

Treatment of *Tmem27* and *BACE2* co-transfected cells with MBi-2 restored levels of full-length TMEM27 to those of cells transfected with *Tmem27* alone, with a concomitant reduction in secreted TMEM27 (Fig. S5). Full-length TMEM27 was not detected in WT islet cell lysates, but was readily detectable in islet lysates from *Bace2* KO mice (Fig. S4c, right). Despite the high level of BACE2 processing of TMEM27 *in vivo*, we were unable to convincingly detect a 25 kDa TMEM27 reactive fragment in the islet culture medium (data not shown). The relative reduction of Aβ 40 in brain cortex and plasma elicited by MBi-1 was comparable across all *Bace2* genotypes and Aβ 40 levels were similar in cortex and plasma in all vehicle-treated animals (Fig. S4a), consistent with BACE2 not contributing to amyloidogenic processing of APP [17]. The difference in the observed MBi-2 IC_50_ value for sAPPβ^swe^ (Fig. S5) compared with previously published value for HEK293 APP^Swe/Lon^ Aβ40 provided in Fig. S2 [49] may be due to differences in the format of the assay (*in vitro* purified enzyme assay vs. whole cell assays) or differences in the substrate.

#### 2.2. *Bace2* KO mice pancreatic islet profiles

BACE2, insulin and glucagon immunohistochemistry in pancreatic islets from WT and *Bace2* KO mice revealed overlap of insulin and BACE2 staining in WT islets, consistent with *Bace2* expression in beta-cells (Fig. S4b). Total islet area was consistent between *Bace2* KO and WT animals, suggesting no difference in islet proliferation between the two genotypes (Table S1).

To examine this more closely, Ki-67-positive cells were counted relative to total nuclei across multiple islets from both *Bace2* KO and WT mice. The percentage of Ki-67-positive cells (∼1%) was consistent between genotypes (Table S1).

#### 2.3. Metabolic assessments in female mice

##### 2.3.1. Response to HFD

The same evident growth delay while on chow diet (CD) associated with *Bace1* ablation seen in males was seen in females (Fig. S6a). Before HFD challenge, both the *Bace1* KO and *Bace1/2* dKO female mice showed highly statistically significant differences in lean mass, consistent with their reduced weights (Fig. S6c); however, there was no statistically significant difference in fat composition for any of the mutant mice as compared with WT by qNMR (Fig. S6c). After HFD challenge, *Bace2* KO females showed no statistically significant difference in total weight, lean mass or fat mass compared with WT, while *Bace*1 KO females had reduced weight and lean mass compared with WT and *Bace*1/2 dKO had reduced weight, lean mass and fat mass compared with WT (Fig. S6d).

##### 2.3.2. Glucose tolerance

No statistically significant differences were noted in OGTT analyses of female mice, other than dKO mice at a single time-point of 20 min (Fig. S7).

##### 2.3.3. Insulin sensitivity

Similarly to males, albeit much less dramatically, genotype effects of consistently depressed blood glucose levels, with improved insulin sensitivity with respect to fasted blood glucose levels for *Bace*1 KO and dKO but not *Bace*2 KO, were seen in females (Fig. S8).

##### 2.3.4. Diet-induced hypercholesterolaemia

For the female mice on CD, total cholesterol and HDL cholesterol were significantly lower (*p* < 0.05) in the dKO mice, and triglycerides and glycerol were higher in the *Bace*1 KO mice (data not shown). No statistically significant differences were detected after HFD challenge in these female mice (data not shown), consistent with their lack of response to HFD challenge.

---

**Table S1.** Characterization of islet density from Bace2 WT and KO mice

| ***Bace2*** | **Islets (n)** | **Average area/islet ($\mu^2$)** |  |  |
| --- | --- | --- | --- | --- |
| WT | 173 | 11,920 |  |  |
| KO | 186 | 10,945 |  |  |
| ***Bace2*** | **Islets (n)** | **Nuclei (n)** | **Ki-67-positive cells (n)** | **Ki-67-positive cells (%)** |
| WT | 23 | 3800 | 39 | 1.03 |
| KO | 9 | 2000 | 21 | 1.05 |

Average area of islets in WT ( $n = 4$ ) vs. KO mice ( $n = 3$ ), and proliferation of beta-cells in WT vs. KO mice, as measured by percentage of Ki-67-positive cells vs. total beta-cell nuclei
*Bace*, beta-site amyloid precursor protein-cleaving enzyme; KO, knockout; WT, wild-type

**Fig. S1.**
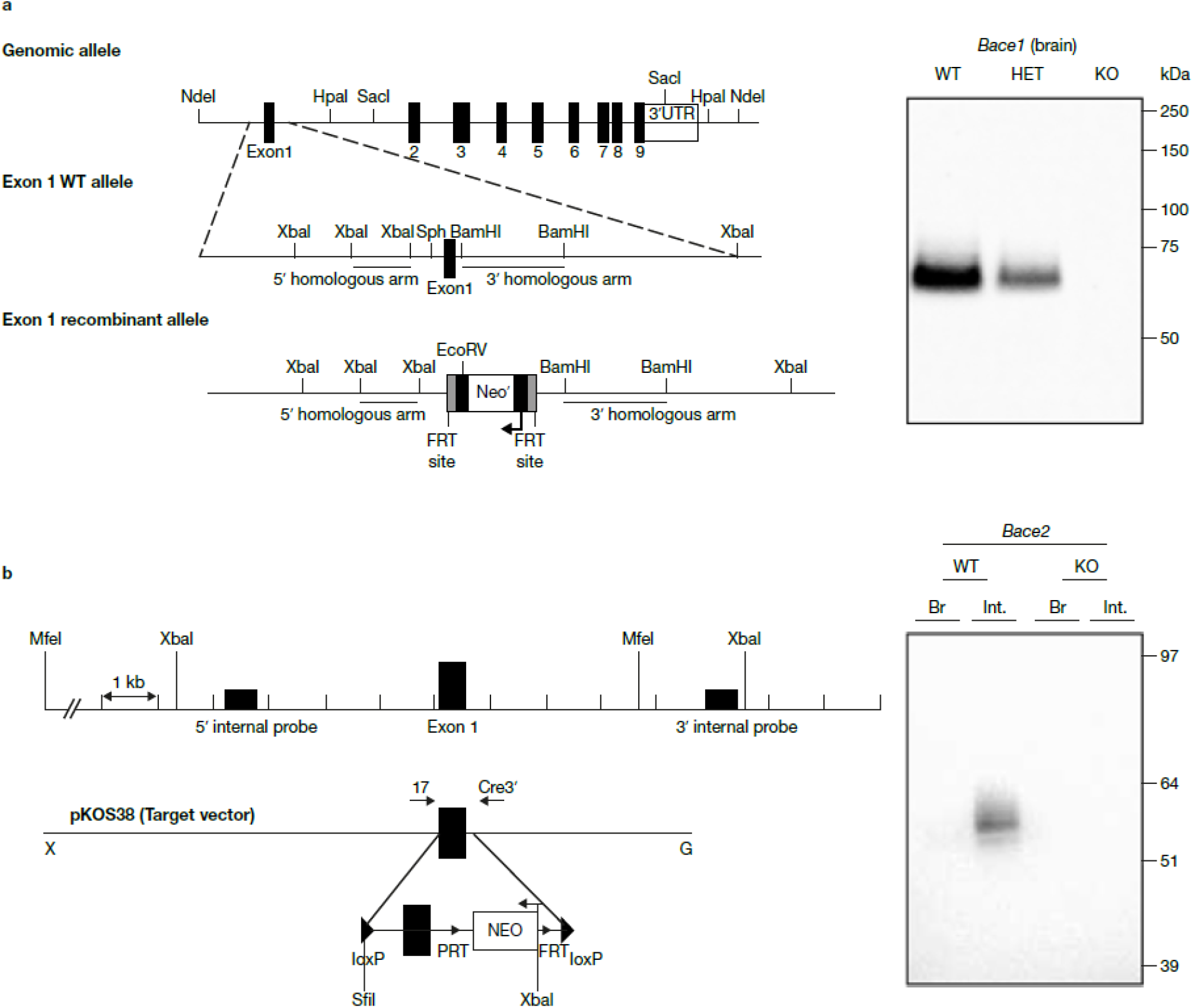
Design and validation of *Bace*1 and *Bace*2 KO mice. (**a**, left) Exon 1-targeting strategy for *Bace1* showing WT and recombinant alleles with polymerase chain reaction primer sequences (underlined). (**a**, right) Western blot of membrane fraction from *Bace1* WT, heterogeneous and KO mouse brains. (**b**, left) Exon 1-targeting strategy for *Bace2*. (**b**, right) Western blot of membrane preparations from 8-week-old *Bace2* WT and KO brain and intestine tissues. Representative images from at least two experiments are shown. *Bace*, beta-site amyloid precursor protein-cleaving enzyme; FRT, flippase recognition target; HET, heterozygous; KO, knockout; UTR, untranslated region; WT, wild-type.

**Fig. S2.**
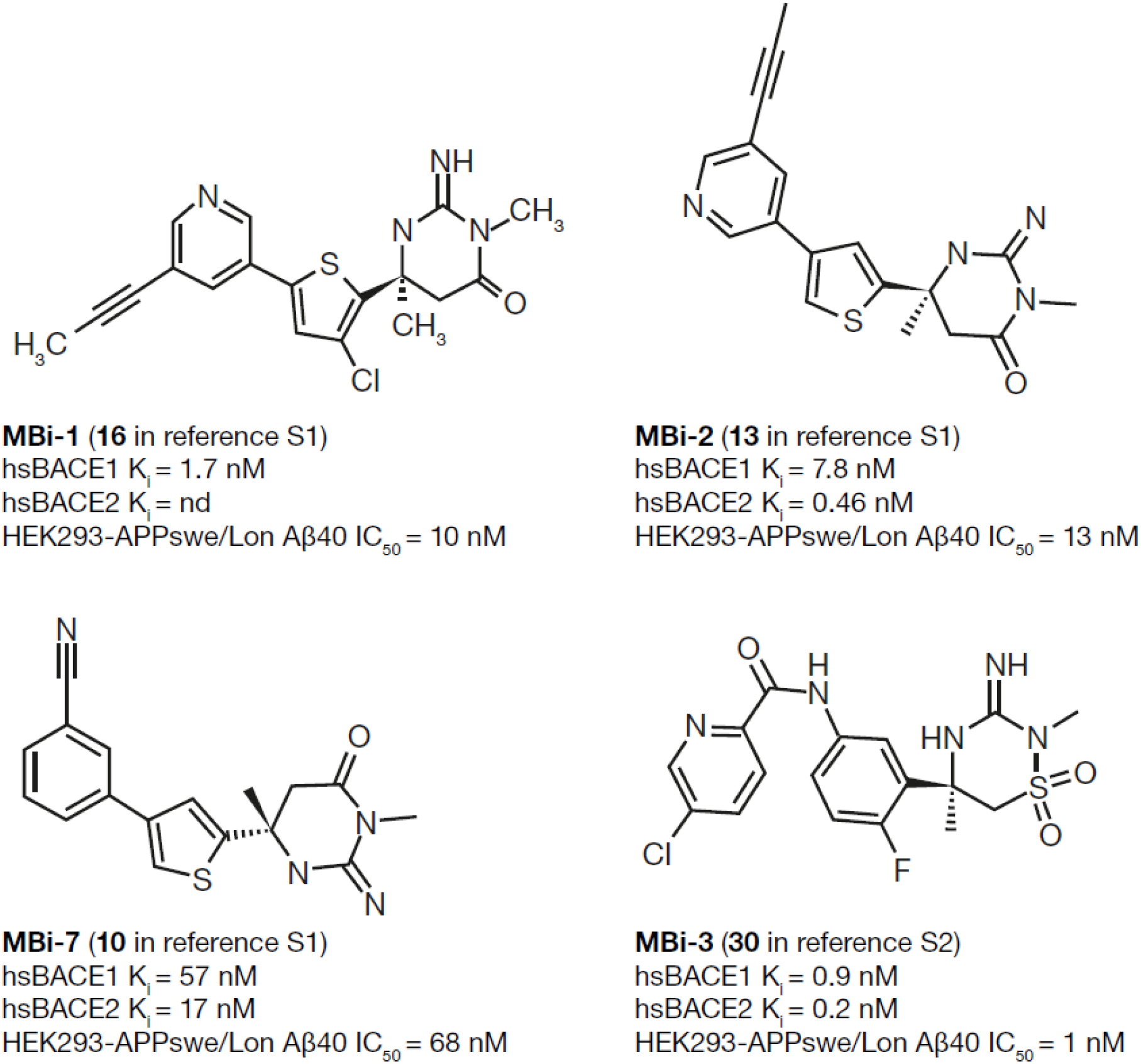
Structures and pharmacological profiles of the BACE inhibitors MBi-1, MBi-2, MBi-7 and MBi-3. See the following references for additional details: [S1] Stamford A et al (2012) ACS Med Chem Lett 3:897-902; [S2] Scott JD et al (2016) J Med Chem 59:10435-10450. *Bace*, beta-site amyloid precursor protein-cleaving enzyme; IC_50_, half-maximal inhibitory concentration; k_i_, inhibition constant, nd, not determined.

**Fig. S3.**
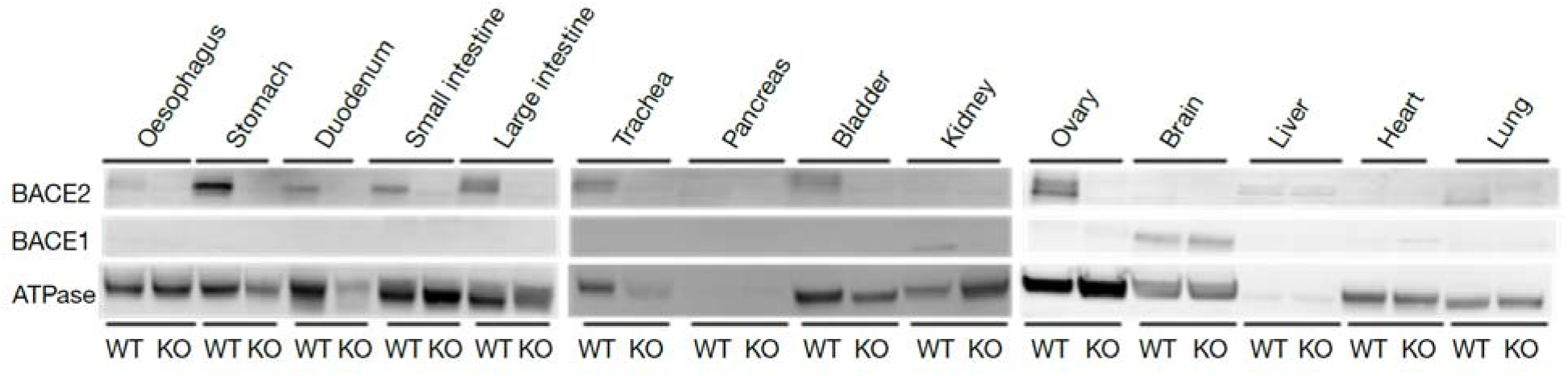
BACE expression profiles across tissues. Western blot profile of BACE1 and BACE2 protein expression in membrane fractions from WT (W) and *Bace2* KO (-/-) mouse tissues. BACE, beta-site amyloid precursor protein-cleaving enzyme; KO, knockout; WT, wild-type.

**Fig. S4.**
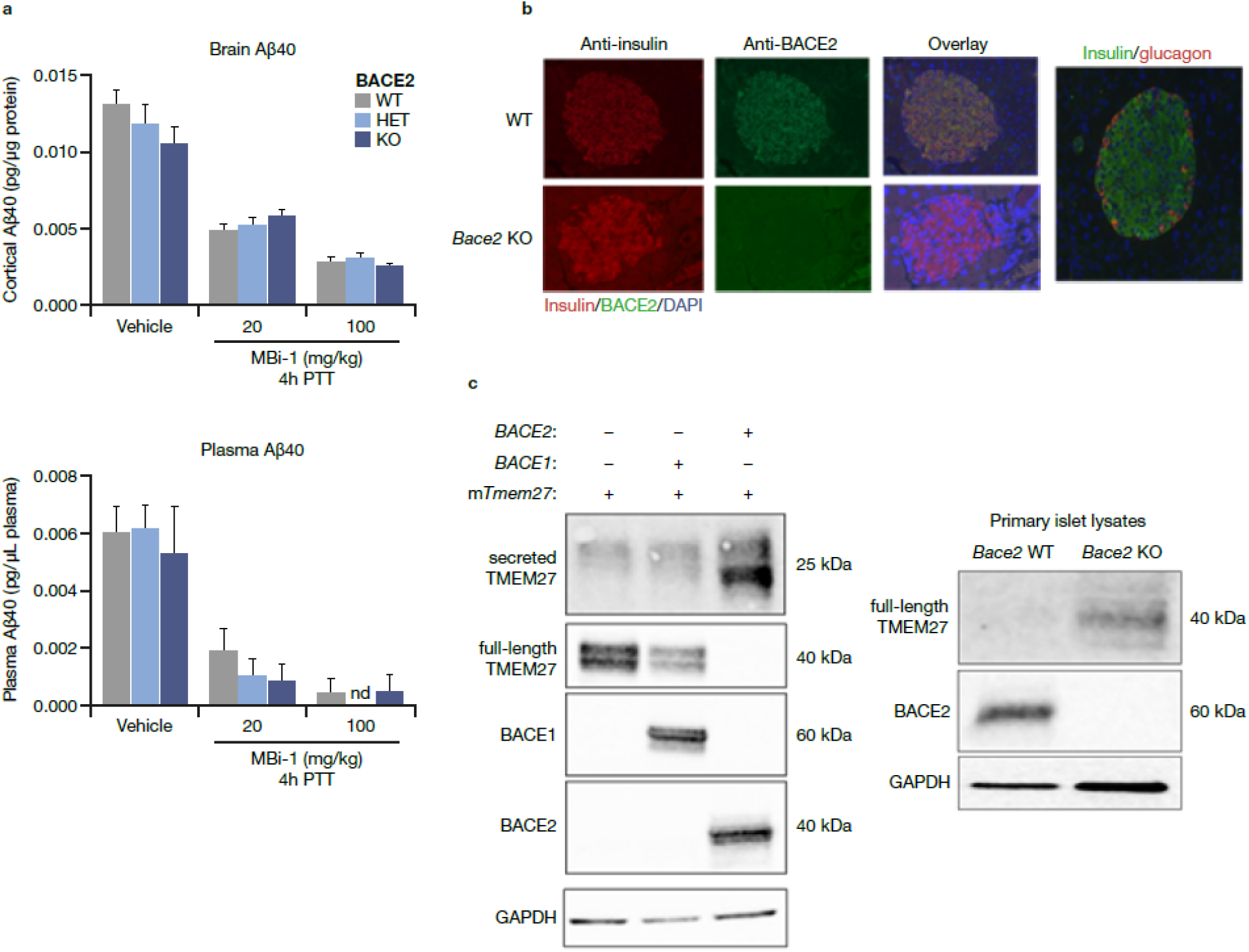
Functional evaluation of *Bace2* WT, heterozygous and KO mice. (**a**) Aβ40 levels in cortex (top) and plasma (bottom) in response to vehicle or a single oral dose of MBi-1 at 20 or 100 mg/kg. (**b**) BACE2, insulin and glucagon immunolocalisation in isolated primary islets from WT but not *Bace2* KO mice. BACE2 immunoreactivity (green) co-localizes with insulin (red) and is consistent with beta-cell expression. DAPI (blue) indicates cell nuclei. (**c**, left panel) HEK293 cells transfected with mouse *Tmem27* with (+) or without (-) co-transfection of *BACE1* or *BACE2*. (**c**, right panel) Primary mouse islets from WT or *Bace2* KO mice. Lysates represent approximately 100 islets. Representative images from at least two experiments are shown. BACE, beta-site amyloid precursor protein-cleaving enzyme; DAPI, 4 ,6-diamidino-2-phenylindole; GAPDH, glyceraldehyde 3-phosphate dehydrogenase; KO, knockout; nd, not detected; PTT, post-treatment time; TMEM27, transmembrane protein 27; VEH, vehicle; WT, wild-type.

**Fig. S5.**
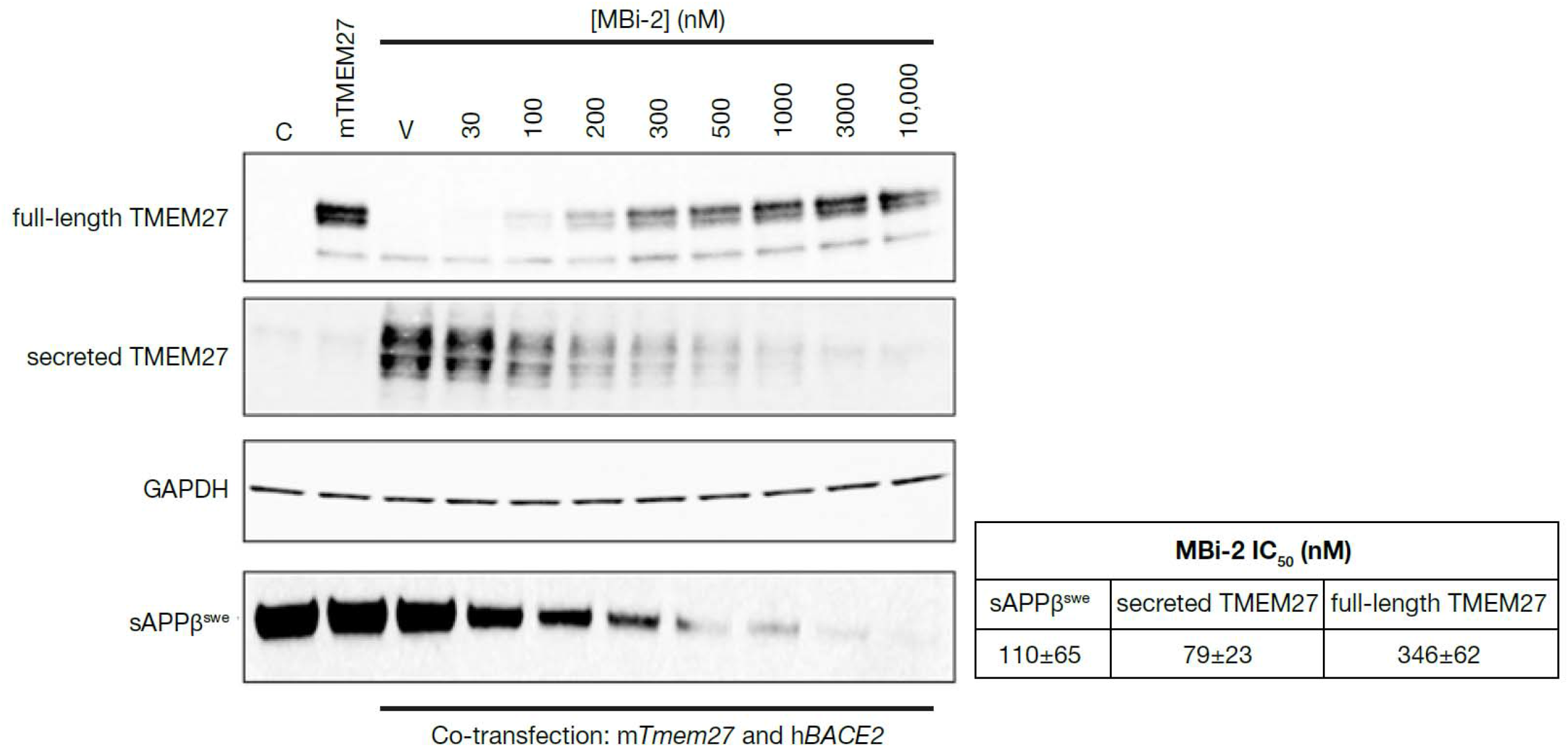
MBi-2 potency for blocking APP vs. TMEM27 processing in transfected HEK293 cells. HEK293-APPswe/lon expressing cells were co-transfected with human *BACE2* and mouse *Tmem-27* followed by treatment with vehicle or increasing concentrations of the BACE inhibitor MBi-2 (see Fig. S2 for MBi-2 structure). C represents vehicle control. The table gives a summary of MBi-2 potency for suppression of sAPPβ^swe^, secreted TMEM27 and elevation of full-length TMEM27. APP, amyloid precursor protein; BACE, beta-site amyloid precursor protein-cleaving enzyme; IC_50_, half-maximal inhibitory concentration; TMEM27, transmembrane protein 27.

**Fig. S6.**
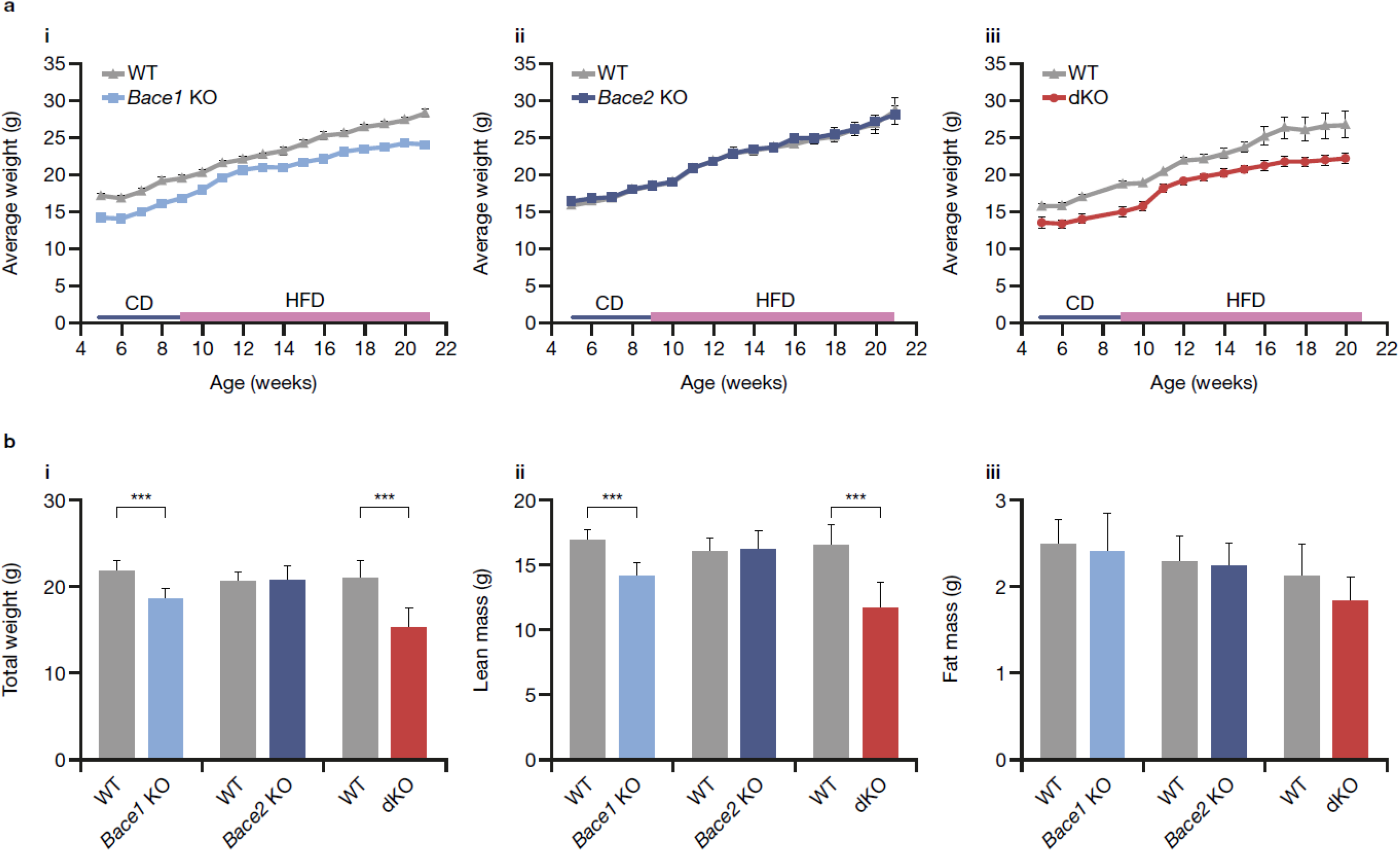
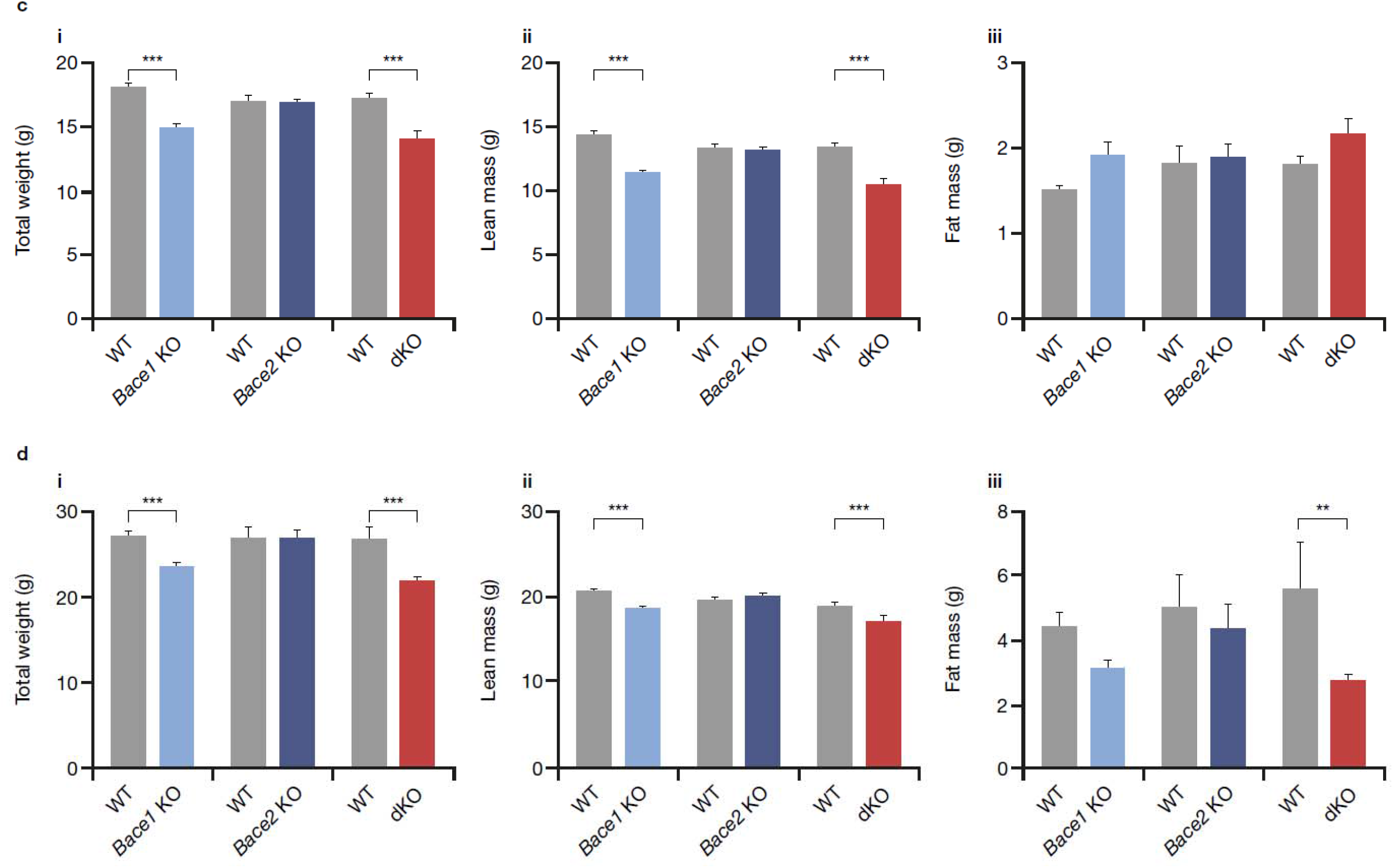
Additional differential response of *Bace* mutants to HFD challenge for growth and body composition. (**a**) Weight-evolution curves of female mice for experiments involving i) *Bace1* KO, ii) *Bace2* KO and iii) dKO mutants. (**b**) Body-composition analysis of male mice before HFD challenge, including i) total weights, ii) lean mass and iii) fat mass. (**c**) Body-composition analysis of female mice before HFD challenge, including i) total weights, ii) lean mass and iii) fat mass. (**d**) Body-composition analysis of female mice after HFD challenge, including i) total weights, ii) lean mass and iii) fat mass. All data are presented as averages ± SD. Statistical analyses performed using Mann–Whitney U test for genotype effect. *\*\*p* < 0.01; *\*\*\*p* < 0.001. *Bace*, beta-site amyloid precursor protein-cleaving enzyme; CD, chow diet; dKO, double *Bace1*/*Bace2* knockout; HFD, high-fat high-cholesterol diet; KO, knockout.

**Fig. S7.**
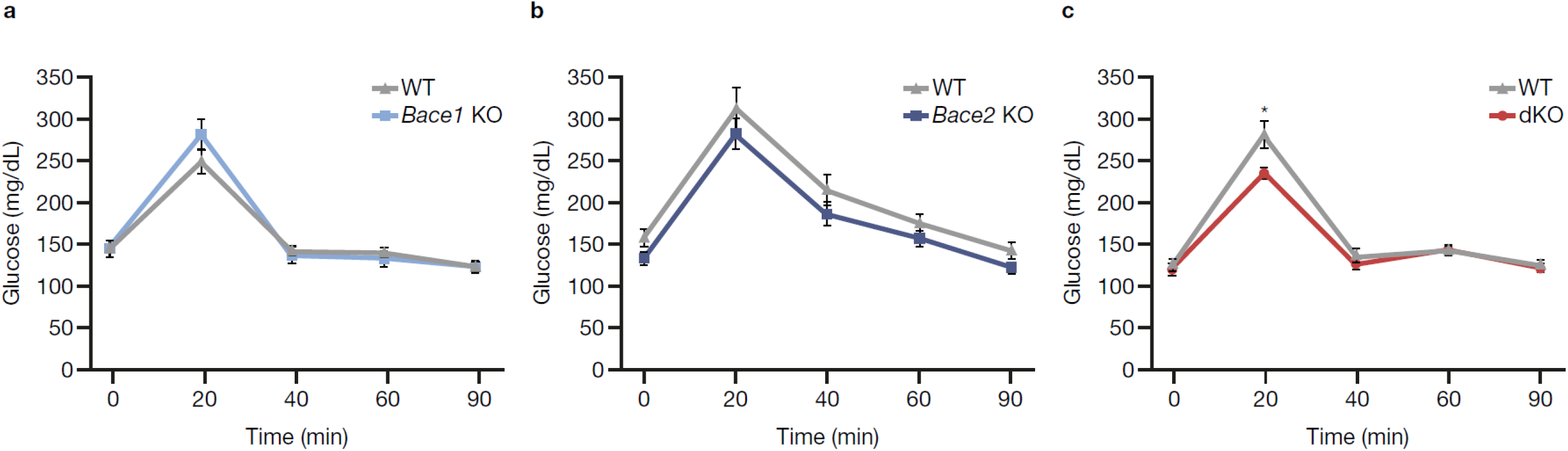
Glucose tolerance testing of *Bace* mutant mice after HFD challenge. Raw blood glucose measurement from fasting (t = 0) and time after glucose bolus (t = 20–90 min) for females of each of the cohorts examined: (**a**) WT vs. *Bace1* KO; (**b**) WT vs. *Bace2* KO; (**c**) WT vs. dKO. All data are presented as averages ± SEM. Statistical analysis performed by Student’s *t*-test for genotype effect. *\*p* < 0.05. *Bace*, beta-site amyloid precursor protein-cleaving enzyme; dKO, double *Bace1*/*Bace2* knockout; HFD, high-fat high-cholesterol diet; WT, wild-type.

**Fig. S8.**
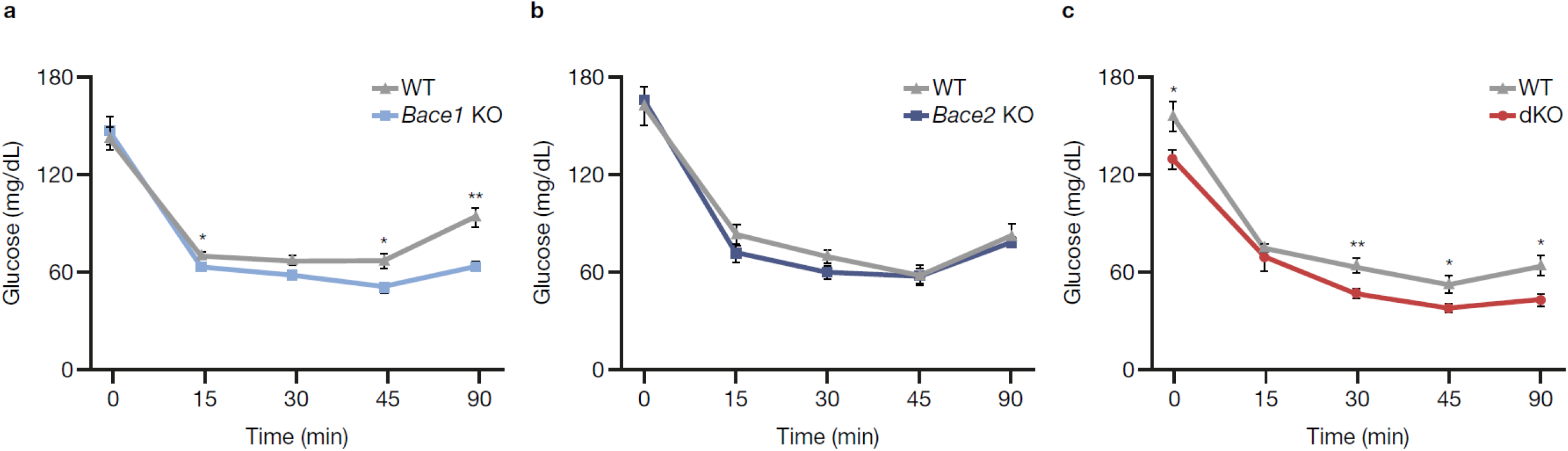
Insulin sensitivity testing of *Bace* mutant mice after HFD challenge. Raw blood glucose measurement from fasting (t = 0) and time after intraperitoneal insulin injection (t = 15–90 min) for females of each of the cohorts examined: (**a**) *Bace1* KO; (**b**) *Bace2* KO; (**c**) dKO. All data are presented as averages ± SEM. Statistical analysis performed by Student’s *t*-test for genotype effect. *\*p* < 0.05;*\*\*p* < 0.01. *Bace*, beta-site amyloid precursor protein-cleaving enzyme; dKO, *Bace1*/*Bace2* double knockout; HFD, high-fat high-cholesterol diet; KO, knockout.

